# C-ToMExO: Learning Cancer Progression Dynamics from Clonal Composition of Tumors

**DOI:** 10.1101/2022.12.23.521788

**Authors:** Mohammadreza Mohaghegh Neyshabouri, Smaragda Dimitrakopoulou, Jens Lagergren

## Abstract

Cancer is an evolutionary process involving the accumulation of somatic mutations in the genome. The tumor’s evolution is known to be highly influenced by specific somatic mutations in so-called cancer driver genes. Cancer progression models are computational tools used to infer the interactions among cancer driver genes by analyzing the pattern of absence/presence of mutations in different tumors of a cohort. In an abundance of subclonal mutations, discarding the heterogeneity of tumors and investigating the interrelations among the driver genes solely based on tumor-level data can result in misleading interpretations. In this paper, we introduce a computational approach to infer cancer progression models from the clone-level data gathered from a cohort of tumors. Our method leverages the rich clone-level data to identify the patterns of interactions among cancer driver genes and produce significantly more robust and reliable cancer progression models. Using a novel efficient Markov Chain Monte Carlo inference algorithm, our method provides outstanding scalability to the rapidly increasing size of available datasets. Using an extensive set of synthetic data experiments, we demonstrate the performance of our inference method in recovering the generative progression models. Finally, we present our analysis of two sub-types of lung cancer using biological multi-regional bulk data.

## Introduction

Cancer progression is known to be an evolutionary process involving the accumulation of somatic mutations [1]. Mutations in specific genes, called cancer driver genes, play critical roles in cancer progression [2]. Comprehending the interactions among mutations in these genes leads to a better overall understanding of the tumorigenesis process, which is crucial for both research and clinical applications. Cancer progression models (CPMs) are tools employed to infer various types of interactions among cancer driver genes, including the temporal order of mutations [3] and the promoting/inhibiting effects they assert on each other [4, 5]. Working with cross-sectional data from cohorts of tumors, the typical setting in CPMs is to evaluate the tumor-level state of a set of genes in each tumor (mutated vs. healthy) and study the cohort-level patterns among the mutations. In this paper, we introduce a method to utilize finer information on the clonal composition of individual tumors for more reliable identification of the interactions among cancer driver genes.

The interactions among cancer driver genes are studied using a developing set of cancer progression models in the literature. A broad set of studies have focused on inferring the temporal order of mutations in the driver genes [6–11]. While randomness is an inherent feature of any evolutionary process, a repetitive temporal order in mutations of specific driver genes is likely to have an underlying biological reason. As an example, a loss-of-function mutation in a tumor suppressor gene 𝒜 might be the key to igniting a specific biological process, increasing the chance of mutation in another driver gene ℬ. In such a scenario, a so-called *progression relation* [12] from gene 𝒜 to gene ℬ is expected to emerge in the observations gathered from a cohort of tumors. In other words, we expect to have an over-representation of mutations in gene ℬ among the tumors that have a mutation in gene 𝒜.

The effects of mutations in the cancer progression process are typically applied through changes in particular biological pathways [13]. As such, mutations in different genes belonging to a specific pathway may exhaust each other’s selective advantage by asserting similar biological effects. Following our previous example, suppose gene 𝒞 plays a similar role as gene 𝒜. In that case, we expect to see a so-called *mutual exclusivity relation* [12] among 𝒞 and 𝒜, meaning that while mutations in any one of these genes 𝒜 or 𝒞 improves the evolutionary fitness of the harboring cells, having a mutation in one of them is enough (in the sense of selective advantage). Moreover, the progression relation from the pathway including both 𝒜 and 𝒞 to gene ℬ might be statistically stronger than the progression relation from 𝒜 to ℬ or from 𝒞 to ℬ. Various methods with different mathematical models and inference techniques are introduced to discover such pathway-level patterns in the driver genes [12–16].

Most cancer progression models (CPMs) defined above are designed to work with tumor-level data on the presence/absence of mutations in specific genes of interest. In this paper, we argue that using such tumor-level data is limiting as the signals are likely to cancel out at such coarse granularity. For instance, continuing with our example above, a single tumor may have mutations in both genes 𝒜 and 𝒞, but in different cells/clones. In this case, this particular tumor would be misleading evidence against the mutual exclusivity of 𝒜 and 𝒞 if we limit our scope to the tumor-level data. Using the tumor-level representation of tumors has been practically inevitable until recently. The rapid developments in data availability and sequencing technologies during recent years provide a more refined understanding of tumor compositions in swiftly growing cohorts of tumors. In particular, we can now investigate the genotype and population of individual clones using single-cell or multi-region bulk data and methods designed to infer the clonal structure from such data. These improvements in the quality of available data further incentivize the development of methods to utilize the clone-level information to provide deeper insight into the cancer progression dynamics.

In this paper, we focus on a recently introduced CPM method called ToMExO [12]. This method uses a probabilistic model for cancer progression. It builds a Markov Chain Monte Carlo (MCMC) algorithm for inferring a *cancer driver tree*, encoding the progression relations among sets of genes or mutually exclusive pathways. As ToMExO provides a good modeling power (of the progression model), together with a computationally efficient likelihood calculation procedure, we assume a ToMExO-like progression model to govern the progression dynamics at the level of individual cells (rather than tumors) and build our model on top of it. Our method can use the clone-level data from the tumors to more reliably identify the mutual exclusivity and progression patterns among the cancer driver genes. We introduce a set of *guided* MCMC moves for making inferences. These novel moves substantially improve the efficiency of our inference algorithm in exploring the space of possible models compared to the random-walk moves used in ToMExO. We call our method C-ToMExO, standing for “Clonal-ToMExO,” as it covers the ToMExO setting as a special case (assuming monoclonal tumors) and extends it to the general case of multi-clonal tumors.

A recently published method called TreeMHN [17] seeks to infer a Mutual Hazard Network (MHN) [5], a type of CPMs, using clonal trees. This method, however, comes with a set of drawbacks that we have addressed in this paper. Firstly, TreeMHN discards the population of the clones, while our method C-ToMExO efficiently uses the population information. We argue that bigger clones are far more important than the small clones, which are also inherently less reliable from the tumor tree reconstruction point of view, i.e., could be misplaced in the tree, leading to wrong genotype inference. TreeMHN is also using a complex progression model, severely limiting its scalability with the increasing number of genes in the data. Our method C-ToMExO, on the other hand, uses a less complex and more interpretable progression model. Our fast and efficient inference algorithm featuring linear computational complexity in both the number of genes and the number of tumors makes it possible to investigate massive datasets.

## 1 Method

The general workflow of our analysis is depicted in Fig. 1. Ultimate goal is to infer the progression dynamics of a cancer type, encoded by the cancer progression model, using cross-sectional data from a cohort of patients. Given a set of samples, preferably from multiple regions of each tumor, we process the data to build a clonal tree for each tumor in the cohort. We then use the clonal trees of all tumors together to infer the general progression model.

**Fig 1.**
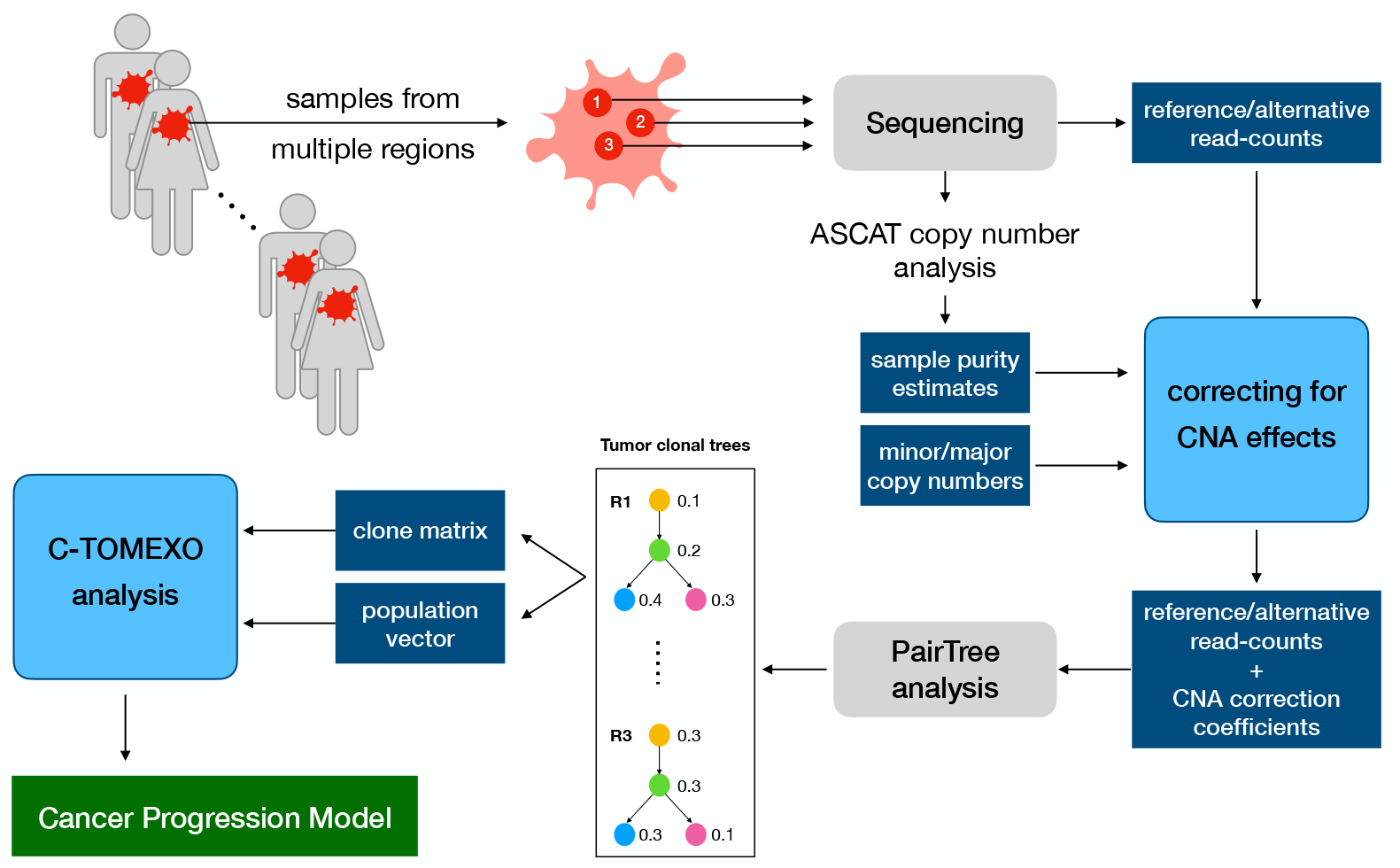
The general workflow of our pipeline for inferring the cancer progression model underlying the tumorigenesis in a cohort of patients. The novel components introduced in this paper are shown using light blue boxes.

**Fig 2.**
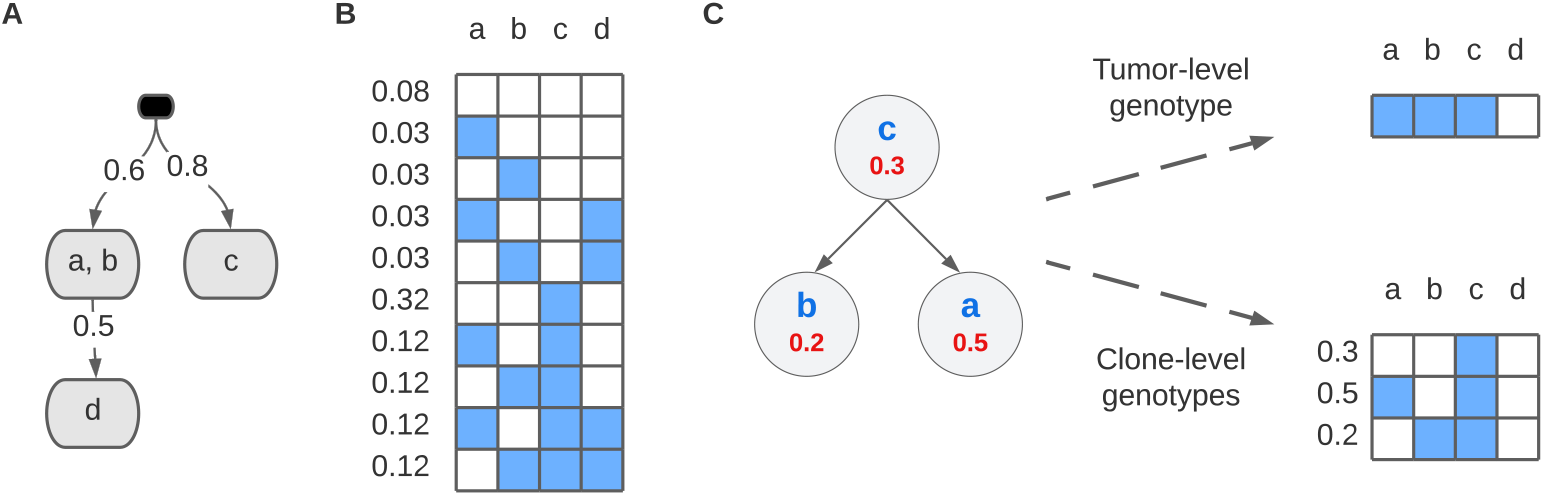
**A**. An example driver tree with four genes **a, b, c**, and **d**. The firing probabilities are shown over the edges. **B**. The implied distribution over the genotypes. Each row represents one possible genotype, with blue elements representing the mutated state, i.e., 1’s. The probability of each row is shown beside it. **C**. An example tumor tree, generated using a phylogeny algorithm from the VAF information. Note that we consider only the genes of interest (driver gene) that are also included in the driver tree. Each node of the tumor tree represents a clone with a relative population shown in red. The clones’ new mutations are shown in blue. Each clone inherits all the mutations in its ancestors and additional mutations in its own node. Two types of representing this tumor by one genotype vector or a tumor matrix together with the population vector are shown on the right-hand side of the figure.

In this section, we start by describing our cancer progression model and how it explains the interrelations among cancer driver genes. We then explain our inference algorithm, C-ToMExO, which is the central novel component of our pipeline. Finally, we describe our novel approach to preprocess the data and correct the effect of copy number alteration (CNA) events on the data. The typical approach for correcting the effect of CNA events is based on assuming that no copy number event happens after the introduction of the somatic mutations. As we explain later in the paper, this assumption leads to many contradictions in our biological data, especially in critical driver genes. Our novel approach for correcting the effect of CNA events relaxes the problematic assumption. It allows for CNA events after the somatic mutations, which is crucial for resolving the seemingly contradictory data and building reliable clonal trees used for the downstream C-ToMExO analysis.

### 1.1 Cancer progression model

In this section, we describe our model for the progression dynamics at the level of individual cells. To this end, we use a probabilistic model previously introduced to model the whole tumors (rather than their constituent cells). We then discuss the drawbacks of assuming such a progression model to govern the behavior of tumors, as it implicitly regards the tumors as homogeneous entities. Finally, we explain our solution for modeling heterogeneous tumors. Our solution allows for exploiting the information on the clonal composition of tumors, if available, to better identify the progression dynamics in cancer.

#### Cancer progression model as a generative process for genotypes

We use a simplified version of the progression model originally introduced in ToMExO [12] as our core model. This model is designed to explain the progression and mutual exclusivity relations among cancer driver genes and is used at the level of tumors in its corresponding publication. In the following, we explain our simplified version of the ToMExO model, which we use at the level of individual cancer cells.

Let us model each cancer cell using a binary vector of length *N*, composed of the state (mutated vs. healthy) of a set of genes *G* = {*g*_1_, …, *g*_*N*_} in the cell. We later refer to this vector as the cell’s genotype vector. Assuming that the healthy cells have no mutation in these driver genes (represented by the genotype being an all-zero vector), the cancer progression model is designed to model the process of accumulation of the somatic mutations. The model is composed of a driver tree (*V, E*) and a set of parameters defined on it. Except for the special empty root node, each node *v* ∈ *V* includes a non-empty subset of the genes in *G*, denoted by *D*_*v*_, such that the set of *D*_*v*_’s form a non-overlapping complete partition of the set of genes *G*, i.e.,

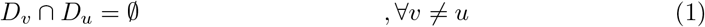

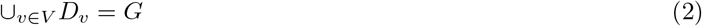

There is a firing probability associated with each *u* → ∈ *v E*. As there is exactly one incoming edge for each node (except the root node), we denote the firing probability of the edge *u* → *v* by *f*_*v*_.

The driver tree explains cancer’s progression dynamics as follows. Starting from the root node, each outgoing edge *u* → *v* ∈ *E* has a chance of firing according to its associated firing probability *f*_*v*_. If the edge *u* → *v* fires, the node *v* gets a mutation in one of its genes (selected according to a categorical distribution). Also, the edges stemming from *v* get their chance to fire; if they fire, the progression process continues along the corresponding edges. Traversing the driver tree following the described process results in a sample genotype, including the state (mutated/healthy) of all genes included in the tree.

A driver tree with *N* genes encodes a distribution over the genotype vectors of length *N*. Consider the example driver tree shown in Fig. 1-A. The genotype distribution implied by this tree is depicted in Fig. 1-B, where the probability of each genotype can be calculated following its generative process. For instance, the probability of having mutation in only gene **a** and gene **c** is the product of the probabilities of the following events:

- The edge pointing to node (**a**,**b**) fires, with probability 0.6,
- Gene **a** gets mutated, with probability 0.5,
- The edge pointing to (**d**) does not fire, with probability 0.5,
- The edge pointing to (**c**) fires, with probability 0.8,

which leads to the overall probability of 0.12 for sampling this genotype. Note that while a total of 2^*N*^ possible genotype vectors of length *N* exist, only a subset of them will have non-zero probabilities in any given driver tree. In our example in Fig. 1-A, only 10 genotypes shown in Fig. 1-B have non-zero probabilities.

#### The necessity of using the clonal representations

As discussed before, CPMs aim to infer a progression model using cross-sectional data from a cohort of tumors. The biological datasets typically consist of called mutations with their number of reference and alternative reads for each tumor. To train any conventional CPM, including ToMExO, we have to summarize the information of each tumor using a single genotype vector. This can be done by, for instance, setting a threshold on the variant allele frequency, such that if the proportion of the variant reads is above a certain threshold, we include the corresponding gene in the set of mutated genes.

In this paper, instead of working with one genotype vector per tumor, we work with a more informative representation of the tumors. As a preprocessing step, we use a method to construct a clonal tree for each tumor using its read-count level information.

Each node of the tumor trees represents one clone. For each clone, we have a set of new mutations, added to those inherited from the parent clone and a relative population of the clone. Let us represent each tumor tree by a binary matrix with the tumor clones in its rows, together with a vector of relative sizes of the clones. We call this representation of the tumor the clone-level genotype representation. We denote the *m*^th^ tumor matrix by *B*_*m*_, where *B*_*m*_[*c*, :] will be the row corresponding to its *c*^th^ clone. Also, we denote the clonal population vector of the *m*^th^ tumor by *θ*_*m*_, where *θ*_*m*_[*c*] is the relative size of the *c*^th^ clone.

Fig 1-C shows an example tumor tree, a tumor-level genotype of the corresponding tumor, and a clone-level genotype representation. This example illustrates the importance of the information we lose in the case of representing the tumor with a single genotype vector. Consider genes **a** and **b**, which are in the same node of the driver tree (see Fig. 1-A) and hence, are expected to be mutually exclusive. Using the variant allele frequency (VAF) information, we could infer (using our tumor tree reconstruction method) that even though we have mutated reads from both genes in this particular tumor, the reads are not from the same subpopulation of the cells. Hence, this tumor in our cohort should not be considered as evidence against the mutual exclusivity of genes **a** and **b**. This misjudgment would be inevitable in the case of representing the tumor using a single vector, as both genes will be considered mutated together.

#### How to use the clone-level representations

Let us assume we take one random cell to represent each tumor. The probability of the cell coming from each clone is proportional to the clone population. Therefore, the genotype of the representative cell will have a categorical distribution over clone genotypes, with the probability of each clone being its relative population.

The method introduced in ToMExO [12] uses a dynamic programming algorithm for calculating the likelihood of observing a genotype, given a driver tree *T* (generating latent unobserved genotypes) and probabilities of false positive *ϵ* and false negative *δ* in the observed genotypes. We emphasize that the model’s ability to handle false positive and false negative events is crucial for analyzing real biological datasets (see [12] for a detailed discussion on this). Using the same algorithm, we can calculate the likelihoods of the form *p*(*B*_*m*_[*c*, :]|*T, ϵ, δ*). We define the likelihood of the *m*^th^ tumor as

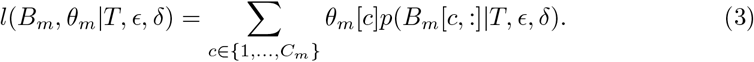

We emphasize that this likelihood can be seen as the expected ToMExO-likelihood of a randomly selected cell from the *m*^th^ tumor. A pseudo-code of our likelihood calculation procedure is included in the Appendix.

### 1.2 C-ToMExO analysis for making inferences

Following a Bayesian inference framework, we want to infer the posterior distribution of the driver tree, given a set of tumors, i.e.,

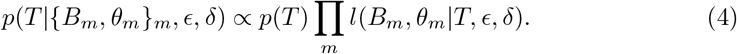

We use a two-step algorithm for making inferences. In the first step, we compute two matrices, the MI matrix (for Mutual Inclusivity) and the ME matrix (for Mutual Exclusivity). These matrices encapsulate the gene-to-gene relations across all clones in the cohort. In the second and main step of the algorithm, we apply a novel, highly efficient Metropolis-Hasting Markov Chain Monte Carlo (MCMC) algorithm to generate samples from the posterior distribution. Our MCMC algorithm incorporates the MI and ME matrices computed in the first step to explore the high probability regions of the space of driver trees in a significantly faster way compared to the ToMExO’s inference algorithm [12].

#### Step 1. Computing the gene-to-gene relation matrices

The space of possible progression models grows exponentially with the number of genes. As a result, an exhaustive exploration of the space quickly becomes infeasible by considering as few as around 20 genes. In this step, we introduce a practical solution for building a *guided MCMC proposal* distribution that enables us to explore the high-probability regions of the progression models space substantially faster than if using the fully random moves used in the ToMExO inference algorithm [12]. This guided proposal distribution is based on the pairwise relationship among the driver genes, encoded by two *N* by *N* matrices (with *N* being the number of genes), with information on the *statistical significance* of the mutual inclusivity (MI) and mutual exclusivity (ME) signals among all pairs of genes.

Discarding the clonal population vectors, for now, we concatenate all the tumor matrices to estimate the overall patterns of mutual exclusivity and mutual inclusivity (co-occurrence) among pairs of genes across the clones. For each pair of genes (*g, w*), let *n*_*g*_, *n*_*w*_ and *n*_*g,w*_ be the number of clones with mutation in *g, w*, and both, respectively. Let *p*_*g*_ be the mutation rate of *g*, i.e., *n*_*g*_ divided by the total number of clones. Using the statistical tests for the mutual exclusivity and the progression relations introduced in the ToMExO paper [12], we can compute the p-value of the promoting or inhibiting effect of mutations in *g* on mutations in *w*, denoted by *p*_PR_(*g, w*) and *p*_ME_(*g, w*), respectively, as

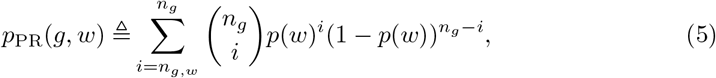

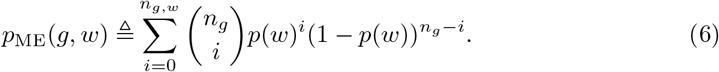

To summarize the significance of mutual inclusivity and mutual exclusivity patterns between the genes, we compute two lower triangular MI and ME matrices, where

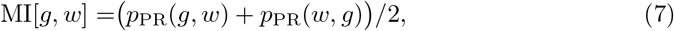

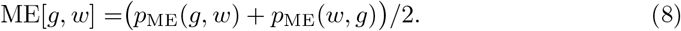

#### Step 2. Guided MCMC algorithm for sampling from the posterior

In this part, we explain the construction of our MCMC algorithm for generating samples from the posterior distribution of the progression models. For the sake of simplicity, we use a uniform prior over the tree structure and the gene assignments, i.e., (*V, E*, {*D*_*v*_}_*v*∈*V*_). In order to restrict our search space, we fix the firing probabilities to a set of empirical estimates, as explained in the following. Consider an edge *u* → *v* from *u* to *v*. To estimate the firing probability of this edge *f*_*v*_, we compute the total population of cells with mutations in the parent node *u* and denote it by 𝒳_*u*_. We then compute the population of cells that have mutations in node *v*, as well as its parent node *u*, and denote it by 𝒴_*v*_. The firing probability of the edge can then be estimated as

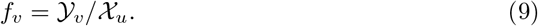

With these firing probability variables, we get a complete prior over the progression models *T* ≜ (*V, E*,{ *D*_*v*_}_*v*∈*V*_, {*f*_*v*_}_*v*∈*V*_) and denote this prior distribution by *p*(*T*).

In order to calculate the likelihood values, we use fixed error probabilities *ϵ* and *δ* in our default setting. As our default error value, we use half of the minimum per-tumor mutation rate among all genes, i.e.,

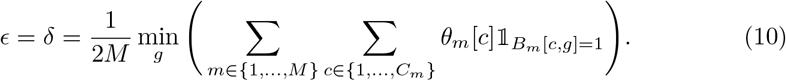

We use a Metropolis-Hasting Markov Chain Monte Carlo (MCMC) algorithm to collect samples from the posterior distribution. As our initial state *T*_0_, we use a star tree structure, where each gene has its node in the first layer. Given the state at round *t, T*_*t*_, we choose from a set of 7 types of moves, shown in Fig. 3, to propose a new candidate state 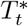. We then calculate our Metropolis-Hasting acceptance ratio as

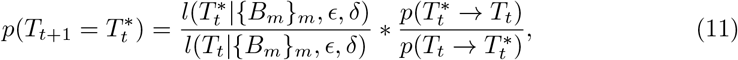

**Fig 3.**
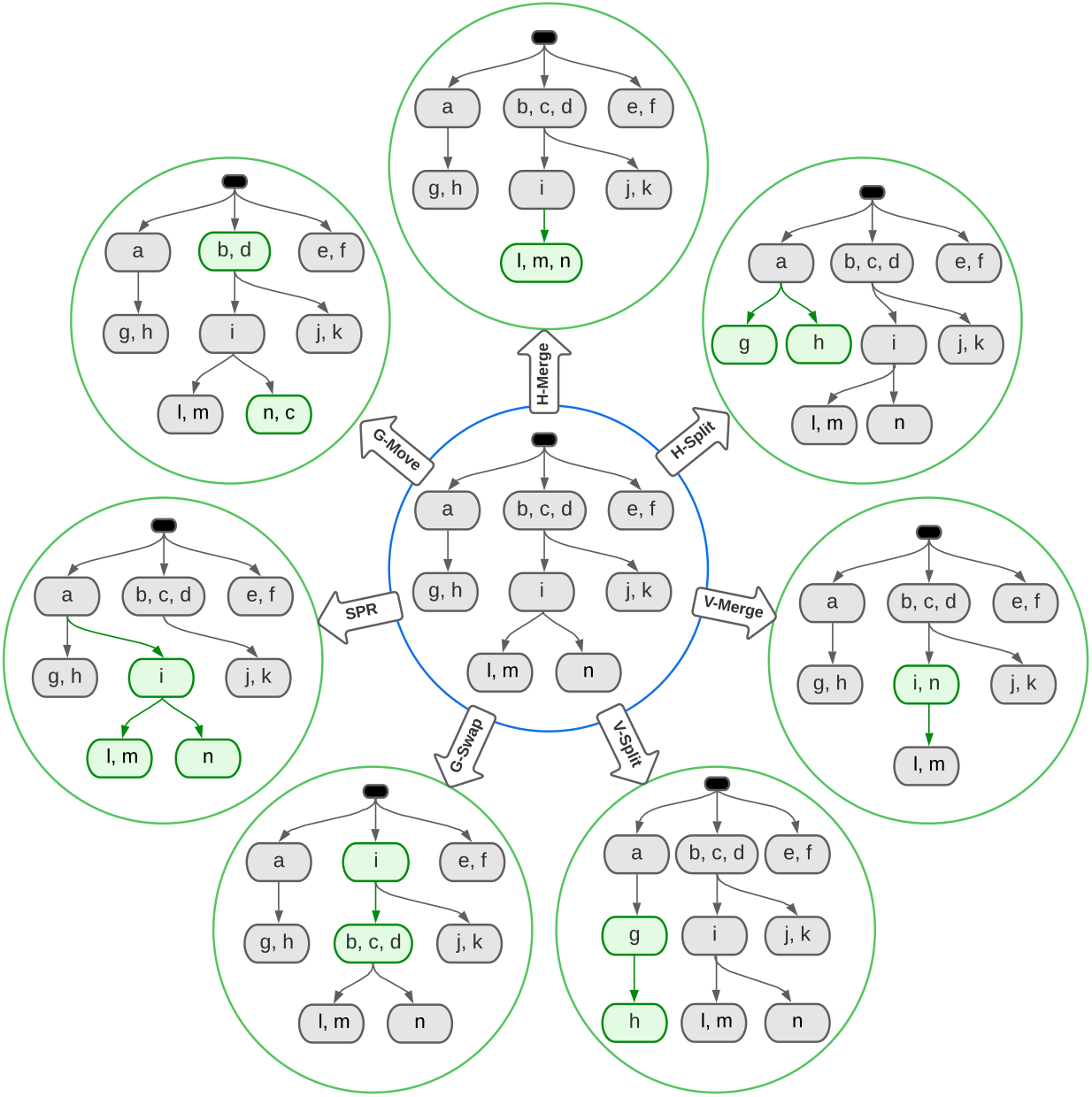
Examples for different types of moves available from any given state of the structure. The driver tree in the center shows the current state of the MCMC, and the driver trees around it inside the green circles are example proposal trees that could arise from different types of available moves. The changing parts of the proposed trees are colored green.

Where 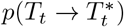 and 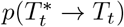 are the forward and backward proposal probabilities that are calculated together with 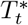 during the proposal procedure. Our structural moves, shown in Fig. 3, include:

1. **V-Merge**: The vertical merge move, where we merge a leaf node into its parent.
2. **V-Split**: The vertical split move, where we detach a subset of a leaf node’s genes and put them into a new child node.
3. **H-Merge**: The horizontal merge move, where we merge a pair of sibling leaf nodes.
4. **H-Split**: The horizontal split move, where we split the set of genes in a leaf node into two subsets and put them into two sibling nodes.
5. **G-Swap**: The gene-swap move, where we swap the genes in a node with the genes in its parent.
6. **SPR**: The Subtree Pruning and Regrafting move, where we detach a node and its subtree and attach it to a new parent.
7. **G-Move**: The single gene move, where a gene is moved from its node into another node.

For each move, the ME and MI matrices calculated in step 1 are used in the following way:

- In splitting moves, nodes with weaker mutual exclusivity among their genes have higher chances of being selected.
- In merging moves, the cases leading to better nodes with stronger mutual exclusivity among their genes have higher chances of being selected.
- In gene-swap moves, nodes with stronger mutual inclusivity with their parents have higher chances of being selected.
- In SPR moves, nodes with stronger mutual inclusivity have a higher chance of getting attached.
- In single gene moves, genes that do not fit in their containing node (with weaker mutual exclusivity with the genes in the same node and weaker mutual inclusivity with the genes in the parent and children nodes) have higher chances of being selected. The destination of a selected gene is also decided with the same criteria for its fitness in the new node.

Using our well-informed proposal distribution leads to a significantly more efficient exploration of the high-probability regions of the posterior distribution. We emphasize that to have proper MCMC mixing and avoid getting stuck in local optimum points, we mix our guided moves with fully random moves, disregarding the MI and ME matrices. In our default setting, we set the probability of fully random moves to 0.5. Moreover, to refrain from overly sharp proposal distributions, we use a softening function to enforce an upper bound equal to 1000 on the ratio of the highest to the lowest probabilities in the categorical distributions we use while constructing samples to propose. The details on our MCMC algorithms can be found in the Appendix.

### 1.3 Correcting the data for the effect of CNA events

One of the critical preprocessing steps for constructing the tumor trees using bulk data is to correct for the effect of copy number alterations (CNAs) in the data (see our pipeline in Fig. 1). In this section, we explain our approach to incorporating the copy number analysis results into our data and prepare the data for the tumor tree reconstruction algorithm.

To reconstruct tumor trees based on bulk read-count data, we need to estimate the cellular prevalence of the mutations from their variant and reference read-count numbers. In the absence of copy number alterations (CNAs) in diploid cells, there are two copies of each genomic position, where only one carries the mutation in the mutated cells. Therefore, the mutation’s variant allele frequency (VAF) is expected to be around half its cellular prevalence. The CNA events complicate the relationship between cellular prevalence and VAF values. In this section, we introduce a model for correcting the effect of the CNA events on the relationship between the cellular prevalence of mutations and their corresponding VAF values.

Consider a multi-regional sample from a tumor. For each region *r*, let *ρ*_*r*_ be the overall sample purity, estimated solely based on a copy number analysis. See Fig. 4 as an example region. There are a total of 7 cells from which 4 are affected by the CNA event. In this case, the CNA-based tumor purity estimate will be *ρ*_*r*_ = 4/7. Let *α*_*m,r*_ and *β*_*m,r*_ be the minor and major copy numbers of the genomic position corresponding to mutation *m* in the CNA-affected cells in region *r*. See Fig. 4 for the *α* and *β* values corresponding to 4 example mutations. Let *v*_*m,r*_ and *t*_*m,r*_ be the number of variant and total reads covering the position of *m* in region *r*. Denoting the cellular prevalence of mutation *m* in region *r* by *ϕ*_*m,r*_, we want to calculate a multiplicative factor *w*_*m,r*_ such

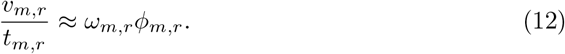

**Fig 4.**
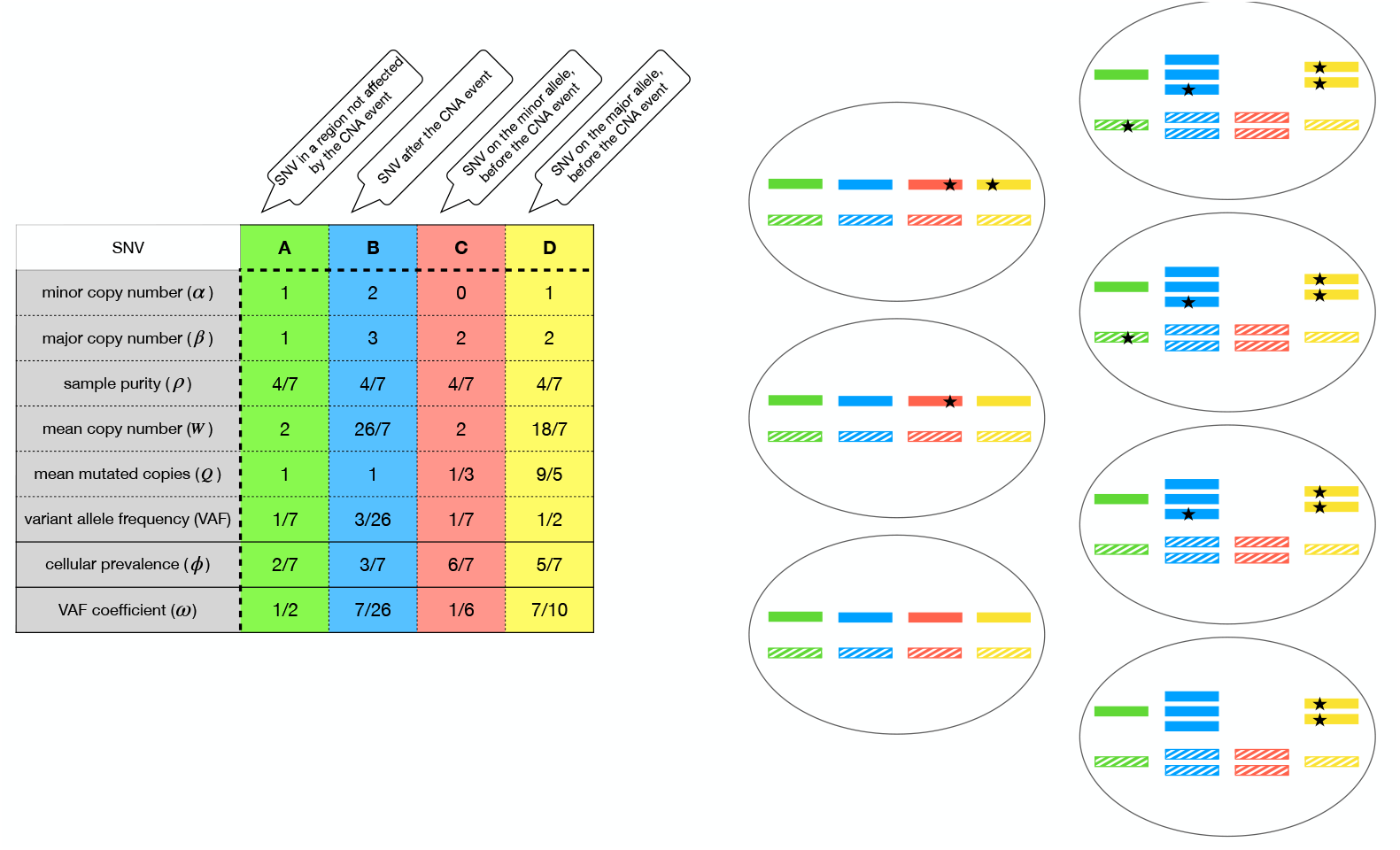
Different scenarios covered by our CNA correction step. There are seven cells in the region shown on the right-hand side. The table on the left shows the variables associated with 4 example mutations. The green and blue mutations (A and B) are of type 1, where the SNV has occurred after the CNA event. The red mutation (C) is of type 2, where the SNV has occurred before the CNA event on the minor allele. In this example, the minor allele is lost due to the CNA event (LOH). The yellow mutation is of type 3, where the SNV has occurred before the CNA on the major allele. In this case, there are some cells with a normal copy number that carry one copy of the mutated allele, while some other cells have two mutated copies.

We emphasize that if the mutation’s position is not affected by the CNA event, i.e., *α*_*m,r*_ = *β*_*m,r*_ = 1, *w*_*m,r*_ will be 1/2. In our example in Fig. 4, mutation *A* (shown in green) represents such a case.

Let *W*_*m,r*_ be the average copy number at position *m* in region *r*. We can calculate *W*_*m,r*_ based on the sample purity and the copy number variables as:

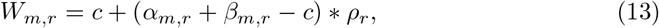

where *c* is the normal copy number, i.e., 1 for males’ sex chromosomes and 2 otherwise. Note that *W*_*m,r*_ is the average number of copies (mutated or healthy) per cell (see Fig. 4 examples). Let *Q*_*m,r*_ be the *average* number of mutated copies in the cells harboring mutation *m* in region *r*. For instance, mutations *A* and *B* in Fig. 4 have *Q* variables equal to 1. Note that multiplying *Q*_*m,r*_ by the cellular prevalence *ϕ*_*m,r*_ will be equal to the average number of mutated copies across all the cells in the region. Therefore, we have

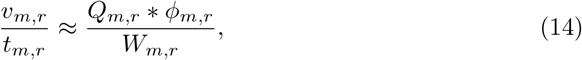

which corresponds to having *w*_*m,r*_ = *Q*_*m,r*_*/W*_*m,r*_.

One typical approach for adjusting *w*_*m,r*_ variables in the presence of CNA events is to assume that all mutations (SNVs) have happened *after* the CNA events. With this assumption, the cells harboring the mutation have only one copy of the mutated allele, i.e., *Q*_*m,r*_ = 1 for all mutations in all regions. This approach for calculating *w*_*m,r*_ variables leads to a wide set of contradictory interpretations of the data when dealing with real biological datasets. With the ultimate objective of constructing the tumor trees, one could discard the *problematic* mutations that lead to contradictions with the assumption of the SNV happening after the CNA events. In practice, such an approach for calculating the *w*_*m,r*_ variables eliminates a broad set of essential driver genes from the tumor trees. However, for our downstream C-ToMExO analysis, we are mainly interested in the relationship between these critical driver genes. Therefore, we cannot afford to discard any of our driver genes while building tumor trees. As such, in the following, we introduce a more concrete way of calculating *Q*_*m,r*_, which allows for more flexibility between the CNA and SNV events. Our approach leads to valid interpretations for all the mutations in the analysis of the biological datasets.

Let’s assume that there is only one copy number event in the history of the tumor evolution, leading to different minor and major copy numbers in different positions. The relationship between each mutation *m* with the CNA event falls into one of the following possibilities:

- **Type 1. SNV after the CNA**: Mutation *m* has happened after the CNA event. In this case, we should have *ϕ*_*m,r*_ ≤ *ρ*_*r*_ for all regions. With no other copy number event, we will have *Q*_*m,r*_ = 1 in this case.
- **Type 2. SNV before the CNA, on the minor allele**: Mutation *m* has happened before the CNA event on the minor allele, which gets *α*_*m,r*_ copies after the CNA event. In this case, *Q*_*m,r*_ depends on the cellular prevalence of the mutation, as the cells without the CNA have only 1 mutated copy, while the cells with the CNA have *α*_*m,r*_ mutated copies:

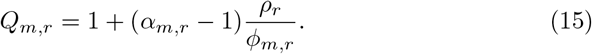 Note that for this case we should have *ϕ*_*m,r*_ ≥ *ρ*_*r*_ for all regions.
- **Type 3. SNV before the CNA, on the major allele**: Mutation *m* has happened before the CNA event on the major allele, which ends up with *β*_*m,r*_ copies after the CNA event. Similar to the previous case, in this case we have:

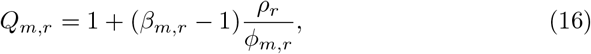

where *ϕ*_*m,r*_ ≥ *ρ*_*r*_ for all regions.

From the example mutations shown in Fig. 4, mutations *A* and *B* are of type 1, mutation *C* is of type 2, and mutation *D* is of type 3.

We follow a Bayesian approach to find the most consistent type for each mutation based on the data collected from all the regions. Since the scenario of type 1 mutations is typically considered the default scenario, we use a prior distribution heavily inclined towards type 1 case, such that except for the mutations on males sex chromosomes, we have:

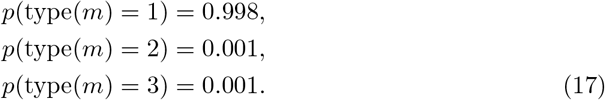

For the mutations on the sex chromosomes in males, as there exists no minor allele, we use the following prior:

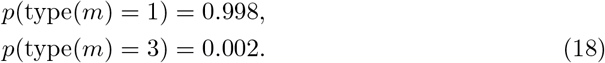

The likelihood of our observations for each mutation *m* can be written as:

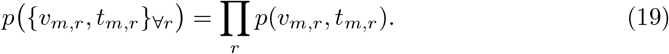

Using a typical binomial model for the read-count data in each region, given the probability of variant read for the position of *m* in region *r* as *ζ*_*m,r*_, we could calculate *p*(*v*_*m,r*_, *t*_*m,r*_) as

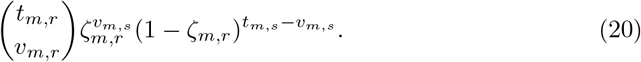

Here, for each one of our three cases, we assume a uniform prior for *ϕ*_*m,r*_ in its valid range and calculate the likelihood of the data given each case by marginalizing out the *ϕ* variable as follows. For type 1 mutations, *ζ*_*m,r*_ = *ϕ*_*m,r*_*/W*_*m,r*_. Hence, we have:

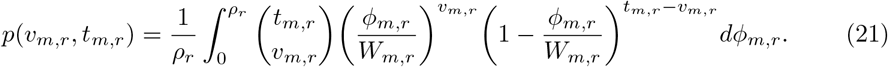

For type 2 mutations, the likelihood can be written as

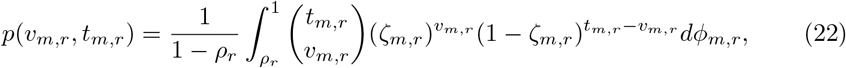

where

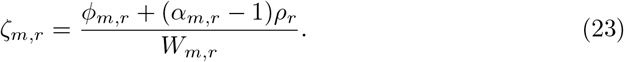

Similarly, for type 3 mutations, we can use Eq.(22) with

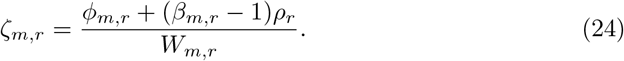

Using the priors and likelihoods described above, we choose the maximum a posteriori type for each mutation. After fixing the mutation type, we calculate the maximum likelihood *Q*_*m,r*_ variables (in their valid ranges according to the mutation types). The *w*_*m,r*_ variables can then be set to *Q*_*m,r*_*/W*_*m,r*_. As the final step, if needed, we adjust the reference counts *r*_*m,r*_ so that Eq. (12) always holds.

## 2 Synthetic Data Experiments

To investigate the performance of our inference algorithm in recovering the generative progression models, we constructed a collection of 100 random generative progression models with:

- the number of genes, from 10 to 40
- the average number of genes per node, from 1 to 4
- the average tree-depth of the genes, from 1 to 4
- the firing probabilities sampled uniformly, from [0.05, 0.95]

With each progression model in the collection, we constructed a set of 12 datasets with the number of tumors in {10, 30, 100, 300} and the error probability in {0.001, 0.01, 0.02}, leading to 1200 synthetic datasets in total. The details of our synthetic data generation procedure can be found in the Appendix.

We used C-ToMExO with a single MCMC chain and 100k iterations to analyze each dataset. Using the maximum a posteriori (MAP) sample as the output of the analysis, we compared it to the known generative model to evaluate our inference algorithm.

In the case of synthetic data experiments, we focus on our success in finding the progression and mutual exclusivity relations implied by the generative driver tree using the F-scores introduced in ToMExO [12]. Fig. 5 shows box plots of our results for different numbers of tumors and error rates. Each scattered circle in this figure represents an experiment. The color of the circles encodes the average tree depth of the genes in the corresponding generative model, with the blue color representing the easier cases with the average tree depth closer to 1, and the red color representing the other side of the spectrum with average tree depths closer to 4. The size of the scattered circles encodes the average number of genes per node in the corresponding generative model, with the cases with more genes per node represented by bigger circles.

**Fig 5.**
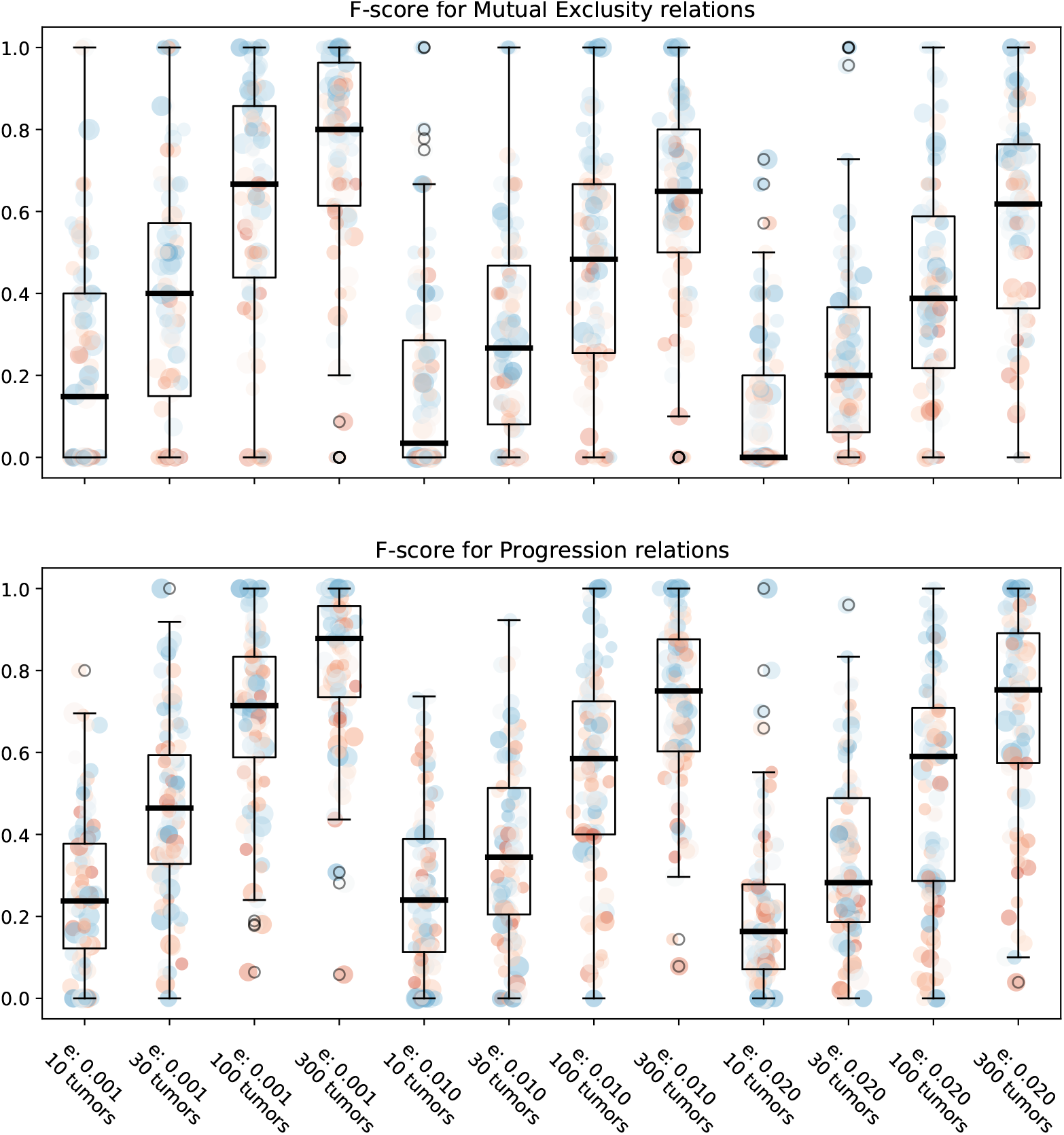
The F-score of C-ToMExO inference algorithm for identification of the mutual exclusivity relationships and progression relationships.

As shown in Fig. 5, a higher number of tumors and a lower noise level leads to more accurate identification of both mutual exclusivity and progression relations, as expected. We emphasize that for the cases with high error rates or few tumors, the generative models are not the best models for explaining the observed data (in the sense of likelihood). As we increase the number of tumors sampled using the generative model, the generative model gradually becomes the most-likely model. In fact, comparing the likelihood of the inferred models to that of the generative models, the inferred models have better likelihoods in almost all cases (1198 out of 1200 experiments). If we use the fixed error parameters used for the inference instead of the generative parameters, the inferred models are still better than the generative models in more than 76% of the experiments (913 out of 1200). These observations show that the inference algorithm is successful in fulfilling its objective of finding the maximum a posteriori (MAP) model.

The F-scores discussed above assume the same importance on all pairwise mutual exclusivity and progression relations. However, some relations are naturally more important in practice. The possible relationship between two driver genes with mutation frequencies of 0.001 among the cancer cells is much more difficult to identify compared to the relationship between two genes with mutation frequencies of 0.5. Moreover, accurate identification of the relationship among highly mutated genes is more important for clinical applications. Note that our progression model puts more emphasis on the highly mutated genes by construction. To have an assessment of how well we identify the mutual exclusivity and progression relations among highly mutated genes in our synthetic data experiments, we define a weighted version of the F-scores, such that the relationship between each pair of genes gets a weight proportional to the product of the genes’ mutation rates. Fig. 6 shows our weighted F-scores. As shown in this figure, for the experiments with as few as 30 tumors, we have a good agreement between the inferred model and the generative one on the relationship they imply on the highly mutated genes. For experiments with 100 or more genes, especially for lower error rates, the inferred model almost totally agrees with the generative one on the important relationships. More details on our synthetic data experiments, including a subset of our random generative progression models and their corresponding parameters, the inferred progression models for the example cases, and the corresponding F-scores, can be found in the Appendix.

**Fig 6.**
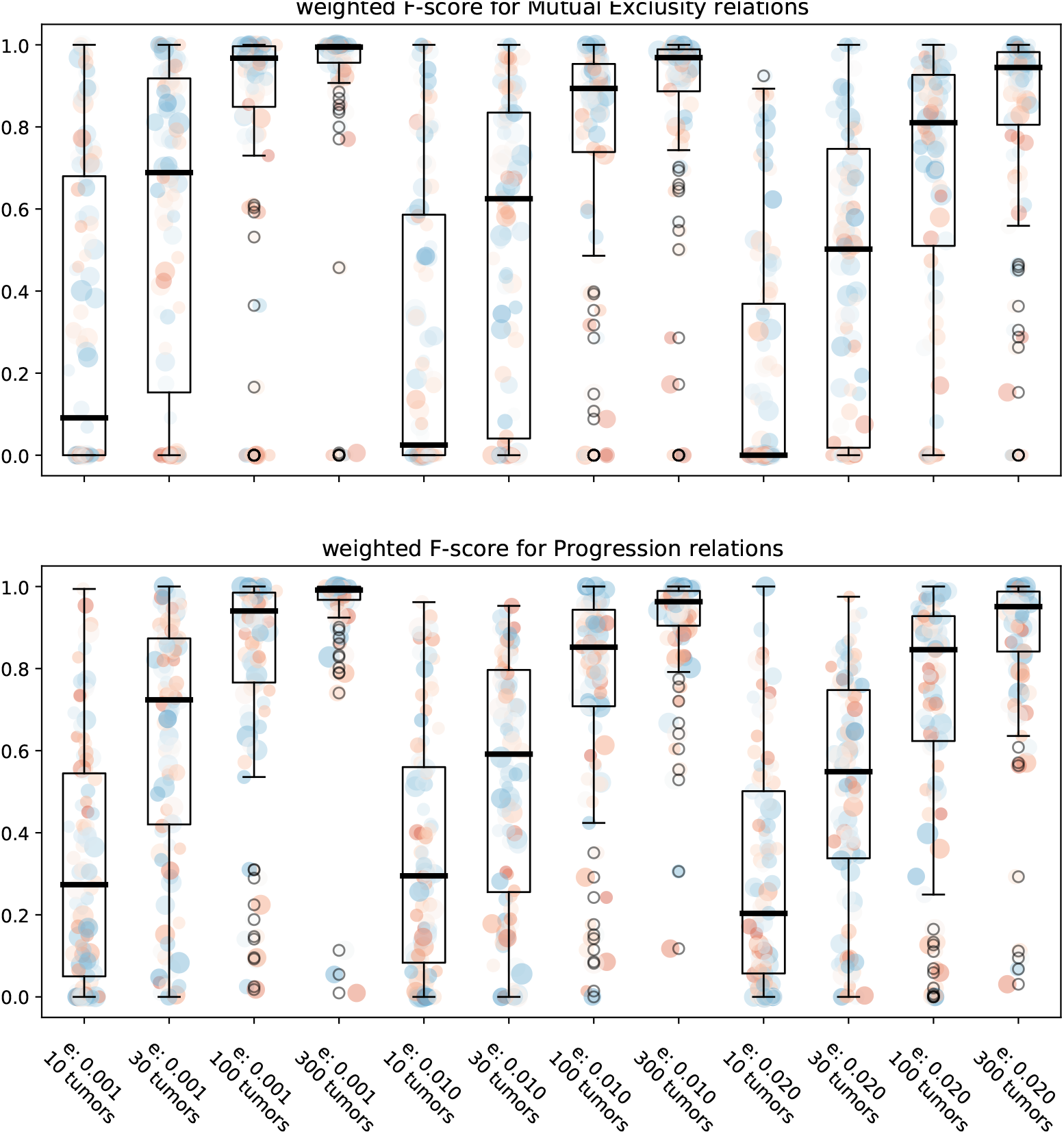
The weighted F-score of C-ToMExO inference algorithm for identification of the mutual exclusivity relationships and progression relationships.

In the following biological data analysis section, we present our results on two biological datasets for LUSC and LUAD cancer types with 32 and 61 tumors, respectively. The error rates we use for the LUSC and LUAD datasets, computed by Eq. (10), are 0.0006 and 0.0018, respectively. Based on our synthetic data experiments, we expect to recover strong mutual exclusivity and progression relations, at least among the highly mutated driver genes.

## 3 Biological Data Analysis

In order to come up with reliable progression models, C-ToMExO heavily relies on the correct reconstruction of the clonal trees. In this paper, we work with the TRACERx Lung cancer dataset [18]. This dataset includes multi-regional bulk DNA sequencing data from a cohort of patients. Samples from various regions of the same tumor can be exploited to improve the accuracy and reliability of the tumor trees significantly. In the following, we start with describing the data. We then review our pipeline for preprocessing the data and preparing our input for C-ToMExO analysis. We finish this section by presenting our inferred progression models and a brief discussion of their biological implications.

### 3.1 TRACERx Lung Cancer dataset

The dataset includes a total of 100 patients. The patients are grouped into 6 subsets based on their cancer subtype. Here, we focus on two sets of invasive adenocarcinoma (LUAD) with 61 patients and squamous cell carcinoma (LUSC) with 32 patients. For each patient, we have a list of 71 to 3706 mutations. As the first step of filtering, we filter out all mutations except splicing and non-synonymous exonic mutations. In the second step of filtering, we only keep the mutations in our candidate driver genes. We use the list of driver genes suggested by IntoGen [2] for lung cancer for this purpose. For each mutation, we have the number of variant reads and total reads for a set of 2 to 7 regions (varying number of regions in different patients). For constructing the tumor trees, we exclude the regions taken from lymph nodes.

The dataset provided by the TRACERx publication [18] includes the results of a copy number estimation analysis using the ASCAT method [19]. The ASCAT analysis results include:

- An estimation of the sample purity for each region, ranging from 0.10 to 0.86.
- An estimation of the major and minor copy numbers for different positions in different regions. Note that the copy number estimation at the same genomic position may vary in different regions of the tumor.

### 3.2 Reconstructing the tumor trees and running C-ToMExO

As shown in our pipeline schematic in Fig. 1, we use PairTree [20] to infer the clonal trees of our tumors. PairTree input is composed of the variant and total read-counts and the multiplicative factors for converting the cellular prevalence to the expected VAF for all mutations and regions. Using a binomial observation model for the counts’ data shown in Eq.(20), PairTree outputs a maximum likelihood estimation of the clonal tree and an estimation of the cellular prevalence of the clones in all regions. Note that the tree structure is the same in different regions, while the cellular prevalence variables may vary from region to region. PairTree has been shown to be able to efficiently leverage the information provided by multi-regional samples to build robust, reliable tumor trees [20]. However, if used without our CNA-correction preprocessing, it discards almost all the driver genes, since the read-count data related to these genes have severe inconsistency with PairTree default assumption on the mutated cells having 1 mutated copy. Discarding the driver genes might not be a deal-breaker for the task of building tumor trees in an abundance of passenger mutations. However, as we specifically need the state of driver genes in the clones, the preprocessing introduced in Section 1.3 preventing PairTree from discarding the driver genes is crucial in our pipeline.

In order to prepare the input for our C-ToMExO analysis, we finally built our tumor matrix from the tree structure, where each row of the matrix holds the genotype of a node in the tree. We took the average population of each clone across all regions as the clone’s population in the tumor. We used 10 MCMC chains with 100k samples for our C-ToMExO analysis for each dataset.

### 3.3 Inferred progression models

We used the TreeMHN method [17] as a baseline to compare against in our analysis of the biological datasets. Fig. 7 shows the connected parts of the Mutual Hazard Networks inferred for the LUAD and LUSC datasets.

**Fig 7.**
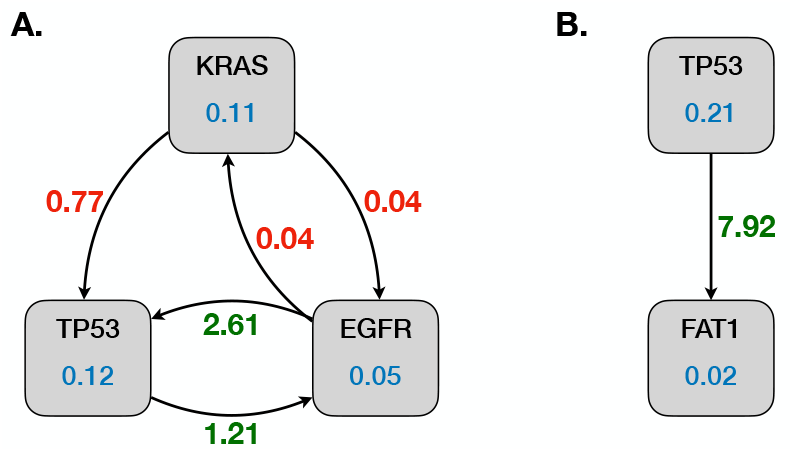
The Mutual Hazard Network (MHN) model inferred by the TreeMHN method for **A**. LUAD dataset, and **B**. LUSC dataset. The mutation rates of the genes are shown in blue inside the nodes. The red/green numbers shown over the edges are the multiplicative inhibiting/promoting effect that mutation in one gene asserts on the other’s mutation rate.

Our inferred progression models for LUAD and LUSC cancer types are shown in Fig. 8 and Fig. 9, respectively. The level of mutual exclusivity and progression relations are shown in blue inside the nodes and over the edges. These scores introduced in [12] take values between 0 and 1, with 1 representing the perfect mutual exclusivity or progression relation. The statistical significance of each signal is shown in green below the corresponding ME or PR score. We emphasize that while our method proposes a complete graph with many implied relations, in contrast with the TreeMHN results, the reported p-values can be used to decide which relationships should be trusted. In the following, we take a closer look at our results.

**Fig 8.**
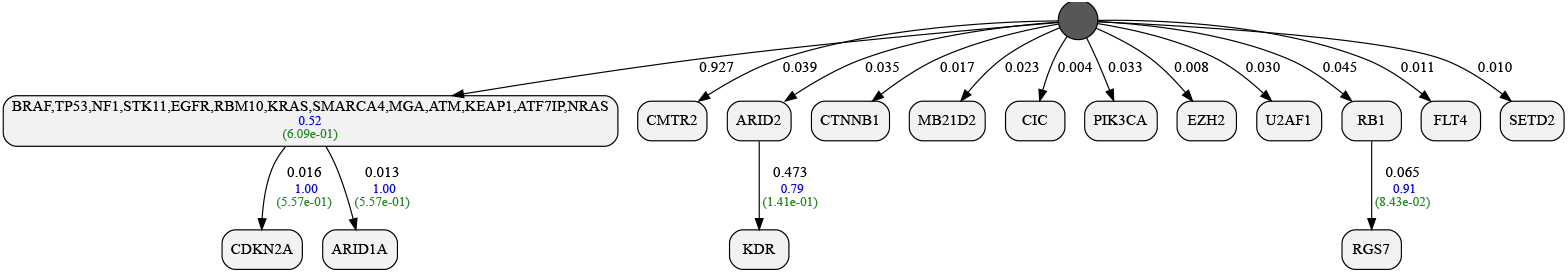
Inferred progression model for Invasive Adenocarcinoma (LUAD) tumors in TRACERx lung cancer data.

**Fig 9.**
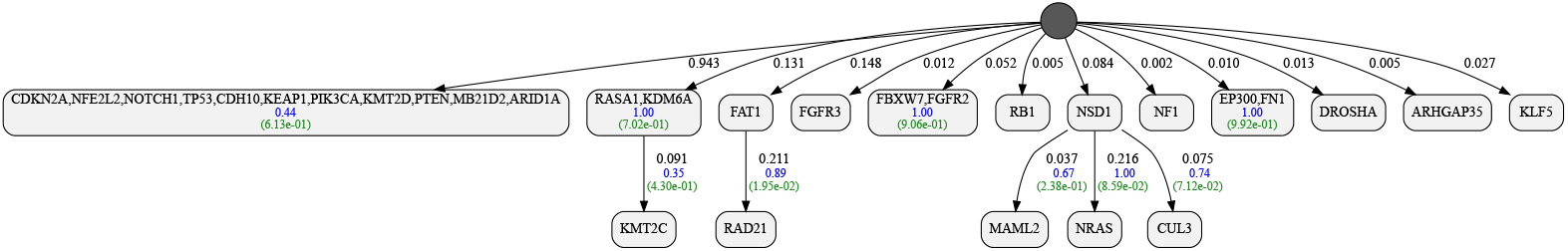
Inferred progression model for Squamous Cell Carcinoma (LUSC) tumors in TRACERx lung cancer data.

#### A. TRACERx LUAD dataset

Fig. 8 shows our inferred progression model for LUAD. Our LUAD dataset includes a total of 61 tumors with 28 genes. The tumor trees we used include between 1 to 6 clones with an average of 2.6 and a standard deviation of 1.4 clones per tumor. The total number of errors in the clones’ genotypes, weighted by their relative population, is 27.3. Our inferred progression model achieves a per-tumor likelihood ratio of 1.88 with respect to the initial star tree, leading to an overall posterior ratio of 4.6 × 10^16^.

Our model for LUAD has EGFR and KRAS together, implying their mutual exclusivity. The mutual exclusivity relationship between mutations in these two genes is well established in the literature with therapeutic implications [21]. There is a quite significant mutual exclusivity relation between these two genes in our dataset with a p-value of 8.5 × 10^−4^, and our inference algorithm has succeeded to discover the relationship. We also have another highly mutated driver gene, TP53, in the same node. TP53 shows slight levels of mutual exclusivity with both EGFR and KRAS in the level of individual clones, which might be interesting for further studies.

#### B. TRACERx LUSC patients

Fig. 9 shows our inferred progression model for LUSC. Our LUSC dataset includes a total of 32 tumors with 30 genes. The tumor trees we used include between 1 to 8 clones with an average of 3.8 and a standard deviation of 1.9 clones per tumor. The total number of errors in the clones’ genotypes, weighted by their relative population, is 24.4. Our inferred progression model achieves a per-tumor likelihood ratio of 2.06 with respect to the initial star tree, leading to an overall posterior ratio of 1.0 × 10^10^.

A comprehensive description of our biological data experiments, including the MI and ME matrices for the datasets and analysis of our inferred posterior distributions, are provided in the Appendix.

## Discussion

In this paper, we introduced a computational method for inferring cancer progression dynamics using cross-sectional data from a cohort of tumors. Our pipeline relies on building clonal trees for the tumors in the cohort and utilizes individual clones for inferring the progression models. We introduced a novel approach for correcting the effect of copy number alterations on the data. This approach leads to a more reliable tumor tree reconstruction, resulting in a more accurate inference of the interrelations among the cancer driver genes. We introduced a guided MCMC algorithm for making inferences, significantly improving our computational efficiency compared to the state-of-the-art. Through an extensive set of synthetic data experiments, we showed the performance of our inference algorithm in various settings. Finally, we analyzed two biological TRACERx lung cancer datasets using our method. With as few as 32 tumors in hand, our method was able to identify a set of interesting relationships among the driver genes. Our synthetic data experiments suggest promising prospects for the applicability of our methods for better identification of the progression dynamics in lung cancer, with more data to be published from the TRACERx cohort in the future.

## Supporting information

S1-Text: method details

## Supporting Information

**S1 Text. Method details.**

## Acknowledgments

This project has received funding from the European Union’s Horizon 2020 research and innovation programme under the Marie Skłodowska-Curie grant agreement

MSCA-ITN-2017-766030 and from the Swedish Foundation for Strategic Research grant BD15-0043.

