## Supplementary material for "C-ToMExO: Learning Cancer Progression Dynamics from Clonal Composition of Tumors": S1-Text: method details

### 1 Model details

#### 1.1 Likelihood calculation procedure

Given a progression model $T = (V, E, \{D_v\}_{v\in V}, \{f_v\}_{v\in V})$, the probability of false positive $\epsilon$, and the probability of false negative $\delta$, expected likelihood of the $m^{\text{th}}$ tumor, represented by a tumor matrix $B_m$ and clonal population matrix $\phi_m$ can be computed using the pseudo-code in Alg. 1.

**Algorithm 1** Calculation of the tumor's expected likelihood

**Require:** $T = (V, E, \{D_v\}_{v\in V}, \{f_v\}_{v\in V})$, $B_m$, $\phi_m$

```
Let α_g = Σ_m Σ_c φ_m[c] B_m[c, g], ∀g
for c ∈ {1, ..., C_m} do                                ▷ C_m: number of clones in the m^th tumor
    for v ∈ post-order(V) do
        o_m[c, v] ← Σ_{g∈D_v} B_m[c, g]
        z_m[c, v] ← |D_v| − o_m[c, v]
        β_m[c, v] ← (Σ_{g∈D_v} B_m[c, g] ∗ α_g)/(Σ_{g∈D_v} α_g)
        Γ_{m,v,c} = ε^{o_m[c,v]} (1 − ε)^{z_m[c,v]}
        Λ_{m,v,c} = β_m[c, v](1 − δ)ε^{o_m[c,v]−1}(1 − ε)^{z_m[c,v]} + (1 − β_m[c, v])δε^{o_m[c,v]}(1 − ε)^{z_m[c,v]−1}
        Ω_{m,v,c} = Γ_{m,v,c} Π_{k∈children(v)} Ω_{m,k,c}
        Ψ_{m,v,c} = Λ_{m,v,c} Π_{k∈children(v)} (f_k Ψ_{m,k,c} + (1 − f_k)Ω_{m,k,c})
    p(B_m[c, :]|T, δ, ε) ← Ψ_{m,root,c}
l(B_m|T, ε, δ) = Σ_{c∈{1,...,C_m}} φ_m[c] p(B_m[c, :]|T, ε, δ)
```

#### 1.2 How to choose the firing probabilities

As explained in the paper, we set the firing probability of each edge $(u, v)$, denoted by $f_v$, to an empirical estimation of the total population of cells with a mutation in both $u$ and $v$, divided by the total population of cells with a mutation in $u$, i.e.,

$$f_v = \mathcal{Y}_v / \mathcal{X}_u, \tag{1}$$

where

$$\mathcal{X}_u = \sum_{m\in\{1,\dots,M\}} \sum_{c\in\{1,\dots,C_m\}} \phi_m[c] \mathbb{1}_{\{\exists g\in D_u | B_m[c,g]=1\}}, \tag{2}$$

$$\mathcal{Y}_v = \sum_{m\in\{1,\dots,M\}} \sum_{c\in\{1,\dots,C_m\}} \phi_m[c] \mathbb{1}_{\{\exists g\in D_v | B_m[c,g]=1\}} * \mathbb{1}_{\{\exists g'\in D_u | B_m[c,g']=1\}}. \tag{3}$$

We emphasize that following the same idea used in ToMExO [2], we could set the firing probabilities to

$$f_v = \max\left\{\frac{\mathcal{Y}_v - \epsilon\mathcal{X}_u}{(1-\epsilon-\delta)\mathcal{X}_u}, 0\right\}. \tag{4}$$

However, as this is proved to be the optimal firing probability only for an exceptional case (when there is only one gene in the dataset), we stick to the more straightforward approach of dividing $\mathcal{Y}_v$ by $\mathcal{X}_v$.

#### 1.3 How to perform error estimation

In our default setting in C-ToMExO, we use a fixed value for the error parameters $\epsilon$ and $\delta$. As our default error value, we use half of the minimum per-tumor mutation rate among all genes, i.e.,

$$\epsilon = \delta = \frac{1}{2M} \min_{g} \left( \sum_{m \in \{1,\ldots,M\}} \sum_{c \in \{1,\ldots,C_m\}} \phi_m[c] \mathbb{1}_{B_m[c,g]=1} \right). \tag{5}$$

We emphasize that our implementation of the method allows for using an empirical error estimation procedure similar to that of ToMExO [2]. Here, we explain this error estimation procedure. We use the dynamic programming algorithm introduced in ToMExO to compute the minimum number of false positives and false negatives for each clone. Building on that, we calculate the expected minimum number of false positives and false negatives in tumor $m$ as:

$$\left(n_m^{FP}, n_m^{FN}\right) = \sum_{c \in \{1,\ldots,C_m\}} \phi_m[c] \left(n_{m,c}^{FP}, n_{m,c}^{FN}\right). \tag{6}$$

Taking a sum over the tumors, we calculate the expected numbers of false positives and false negatives:

$$\left(n_{FP}, n_{FN}\right) = \sum_{m \in \{1,\ldots,M\}} \left(n_m^{FP}, n_m^{FN}\right) \tag{7}$$

Our estimate of $\epsilon$ and $\delta$ will be

$$\left(\hat{\epsilon}, \hat{\delta}\right) = \left(\frac{n_{FP}}{\mathcal{Z} - n_{FN} + n_{FP}}, \frac{n_{FN}}{\mathcal{O} - n_{FP} + n_{FN}}\right), \tag{8}$$

where $\mathcal{O}$ and $\mathcal{Z}$ are the expected number of ones and zeros, i.e.,

$$\mathcal{Z} = \sum_{m \in \{1,\ldots,M\}} \sum_{c \in \{1,\ldots,C_m\}} \phi_m[c] \sum_i \mathbb{1}_{B_m[c,i]=0},$$
$$\mathcal{O} = \sum_{m \in \{1,\ldots,M\}} \sum_{c \in \{1,\ldots,C_m\}} \phi_m[c] \sum_i \mathbb{1}_{B_m[c,i]=1}.$$

Note that the error parameters can also be fine-tuned with a Gradient Ascent algorithm, using a straightforward extension of the formulas used in ToMExO [2].

#### 2 Inference algorithm details

##### 2.1 Smoothing the proposal distribution

As explained in the paper, we use a softening function to refrain from having overly sharp probabilities when selecting from a list of candidates. We use *s-softmax* function, defined in the following, for this purpose.

Let $W = (w_1, \ldots, w_n)$ be a vector of un-normalized log-scaled weights we want to use for drawing a sample, such that the probability of selecting the $i^{\text{th}}$ candidate is proportional to $e^{w_i}$. Let $w_{\min}$ and $w_{\max}$ be the minimum and maximum values in $W$. In order to enforce an upper-bound of $\rho = 1000$ on $e^{w_{\max} - w_{\min}}$, we define:

$$s \triangleq \min\left\{1, \frac{\log \rho}{w_{\max} - w_{\min}}\right\}. \tag{9}$$

Probability of selecting the $i^{\text{th}}$ candidate is then calculated as:

$$p(i) = \frac{e^{sw_i}}{\sum_k e^{sw_k}}. \tag{10}$$

##### 2.2 MCMC proposal distribution

In this section, we go over the details of our proposal distribution. We assume that the ME and MI matrices are available, and we want to have a guided move, taking advantage of these matrices with a probability equal to $\zeta = 0.5$. As the first step, we randomly select a move type. In our default setting, we have the probability of selecting **H-Merge**, **H-Split**, **V-Merge**, **V-Split** and **G-Swap** equal to 1/15, and the probability of **SPR** and **G-Move** equal to 1/3. In the following, we explain the guided type of each move. Note that the random (unguided) moves are identical to the guided moves, except for all the categorical distributions we compute in the process that should be replaced by uniform categorical distributions.

- **H-Merge**:
  - Form a set of candidates of form $\{u, v\}$, where $u$ and $v$ are sibling leaf nodes.
  - Form weights vector $W$ such that

$$W_{\{u,v\}} = -\log \frac{\sum_{g \in D_u} \sum_{w \in D_v} \text{ME}[g, w]}{|D_u| * |D_v|} \tag{11}$$

  - Select a candidate pair of nodes to merge based on s-softmax($W$)

- **H-Split**:
  - Form a set of candidate leaf nodes $u$, where $|D_u| > 1$.
  - Form weights vector $W$ such that

$$W_u = \log \frac{\sum_{\{g,w\} \subset D_u} \text{ME}[g, w]}{\binom{|D_u|}{2}} \tag{12}$$

  - Select a candidate leaf node $v$ to split based on s-softmax($W$)

- Partition $D_v$ into two non-empty subsets $D_{v1}$ and $D_{v2}$, using a uniform distribution (over valid partitions) and form two sibling leaves $v1$ and $v2$.

- **V-Merge**:
  - Form a set of candidate leaf nodes $u$, such that parent of $u$, $\text{pa}(u)$, is not the root node.
  - Form weights vector $W$ such that

$$W_u = -\log \frac{\sum_{g \in D_u} \sum_{w \in D_{\text{pa}(u)}} \text{ME}[g, w]}{|D_u| * |D_{\text{pa}(u)}|} \tag{13}$$

  - Select a candidate node to merge into its parent based on s-softmax($W$)

- **V-Split**:
  - Form a set of candidate nodes $u$, where $|D_u| > 1$.
  - Form weights vector $W$ such that

$$W_u = \log \frac{\sum_{\{g,w\} \subset D_u} \text{ME}[g, w]}{\binom{|D_u|}{2}} \tag{14}$$

  - Select a candidate leaf node $v$ to split based on s-softmax($W$)
  - Select a non-empty non-full subset of $D_v$ to be replaced into a new child of $v$, using a uniform distribution over valid subsets $\{G \subset D_v : |G| > 0, |D_v \setminus G| > 0\}$ and form the new node child.

- **G-Swap**:
  - Form a set of candidate nodes $u$, such that parent of $u$, $\text{pa}(u)$, is not the root node.
  - Form weights vector $W$ such that

$$W_u = -\log \frac{\sum_{g \in D_u} \sum_{w \in D_{\text{pa}(u)}} \text{MI}[g, w]}{|D_u| * |D_{\text{pa}(u)}|} \tag{15}$$

  - Select a candidate node $v$ based on s-softmax($W$) and swap the genes in $D_v$ with the genes in $D_{\text{pa}(u)}$.

- **SPR**:
  - Select a node $u$ using a uniform distribution over all nodes (except the root).
  - Form a set of candidate new parent nodes $v$, such that $v$ is not among the descendants of $u$.
  - Form weights vector $W$ such that $W_{\text{root}}$ is the average of $\log \text{MI}$ matrix (taking the average *after* taking the logarithm). For other nodes in the set of candidates:

$$W_v = -\log \frac{\sum_{g \in D_u} \sum_{w \in D_v} \text{MI}[g, w]}{|D_u| * |D_v|} \tag{16}$$

- Select a candidate node $k$ based on s-softmax($W$) and set $k$ as the new parent of $u$.

- **G-Move**:
  - Form a set of candidate genes $g$ from the nodes that contain at least 2 genes.
  - Let $N(g)$ denote the node containing $g$. Calculate $W_{\mathrm{ME}}$ as

$$W_{\mathrm{ME}} = \log \frac{\sum_{w \neq g \in D_{N(g)}} \mathrm{ME}[g, w]}{|D_{N(g)}| - 1}. \tag{17}$$

  - If $\mathrm{pa}(N(g))$ is the root node, let $W_{\mathrm{MI,p}}$ be the average of $\log \mathrm{MI}$ matrix (taking the average *after* taking the logarithm). Else, calculate

$$W_{\mathrm{MI,p}} = \log \frac{\sum_{w \in D_{\mathrm{pa}(N(g))}} \mathrm{MI}[g, w]}{|D_{\mathrm{pa}(N(g))}|}. \tag{18}$$

  - If $N(g)$ has no children, let $W_{\mathrm{MI,c}}$ be the average of $\log \mathrm{MI}$ matrix (taking the average *after* taking the logarithm). Else, calculate

$$W_{\mathrm{MI,c}} = \log \frac{\sum_{w : N(g) = \mathrm{pa}(N(w))} \mathrm{MI}[g, w]}{|\{w : N(g) = \mathrm{pa}(N(w))\}|}. \tag{19}$$

  - Form weights vector $W$ as

$$W = W_{\mathrm{ME}} + W_{\mathrm{MI,p}} + W_{\mathrm{MI,c}} \tag{20}$$

  - Select a gene $l$ to move based on s-softmax($W$).
  - Form a set of candidate destination nodes $u$ (including all the nodes except the root node).
  - For each node $u$ in the candidate destinations, calculate $W'$ similar to Eq. (20), assuming $l$ is moved into $u$.
  - Select a destination node for $l$ based on s-softmax($-W'$).

We emphasize that in all the moves, we keep track of the forward and backward probabilities to be used in calculating the Metropolis-Hasting acceptance ratio, as explained in the main manuscript.

### 3 Details on the synthetic data experiments

#### 3.1 Sampling tumors from driver trees

In our framework, each progression model imposes a distribution over the binary genotype vectors. Following the generative process described in the paper we can sample genotypes from any given progression model, where each genotype represents a single clone. In this section, we explain a way to modify our genotype sampling procedure into a two-step process. In the first step, we sample a random distribution over a set of genotypes, according to which we sample a random genotype in the second step. In our synthetic data experiments, we take the distribution constructed in the first step as a simulated tumor. We emphasize that this approach perfectly fits with our interpretation of tumors in the C-ToMExO model.

The generative process for sampling random genotype vectors consists of a set of *hard* decisions that are needed to be taken in a pre-order traversal of the driver tree. These hard decisions include decisions on whether specific edges fire or not and decisions on which gene to mutate in a mutated node. Of course, the hard decisions taken during the generative process affect the questions encountered later on. For instance, if we have processed the chance of mutation in a node $u$ and it is not mutated, we won't need to see whether its outgoing edges fire or not.

The key to reforming this sampling procedure into a two-step process is to postpone some of the hard decisions until after traversing the tree. Suppose we are given a set of edges and nodes in which we want to make *soft* (fuzzy) decisions. After a full traversal of the tree, we get a distribution over a subset of the binary vectors. During the traversal, when we face a soft decision point, we cover all possible scenarios with some randomized weights as follows:

- In order to make a soft decision over an edge $(u, v)$ with firing probability $f_v$, we draw a sample $x \sim \text{Beta}(\alpha f_v, \alpha(1 - f_v))$, where $\alpha$ is an arbitrary parameter (we use 10 as a default value). We multiply the weights of the vectors stopping at the current node by $1 - x$. The weights of the vectors that will progress along the edge get multiplied by $x$.

- In order to make a soft decision for selecting a gene among $\{g_1, \ldots, g_n\}$, we draw a sample $x \in \mathbb{R}^n \sim \text{Dirichlet}(\beta * \mathbb{1})$, where $\beta$ is an arbitrary parameter and $\mathbb{1}$ is the vector of ones. We use 10 as a default value for $\beta$. We multiply the weights of the vectors adding mutation in $g_i$ by $x_i$ and proceed with the corresponding vectors independently. We could also use the same continuation process for all these vectors with different $g_i$'s.

In the second step, we draw a sample from the distribution resulting from the first step and output the sampled binary vector.

While any edge or node can be considered for soft decisions, we choose the soft decision set in a particular manner:

- We randomly select a leaf node,

- We select a random subset of the edges in the path from the selected leaf to the node for soft decision.

Such a set of soft decision edges results in a distribution over binary vectors that resembles a dataset derived from a linear tumor tree. Therefore, we can use the output of the first step as a simulated tumor. Fig. 1 shows an example progression model and two tumors sampled from it. The process of building the matrices as we go down the progression model is shown for better

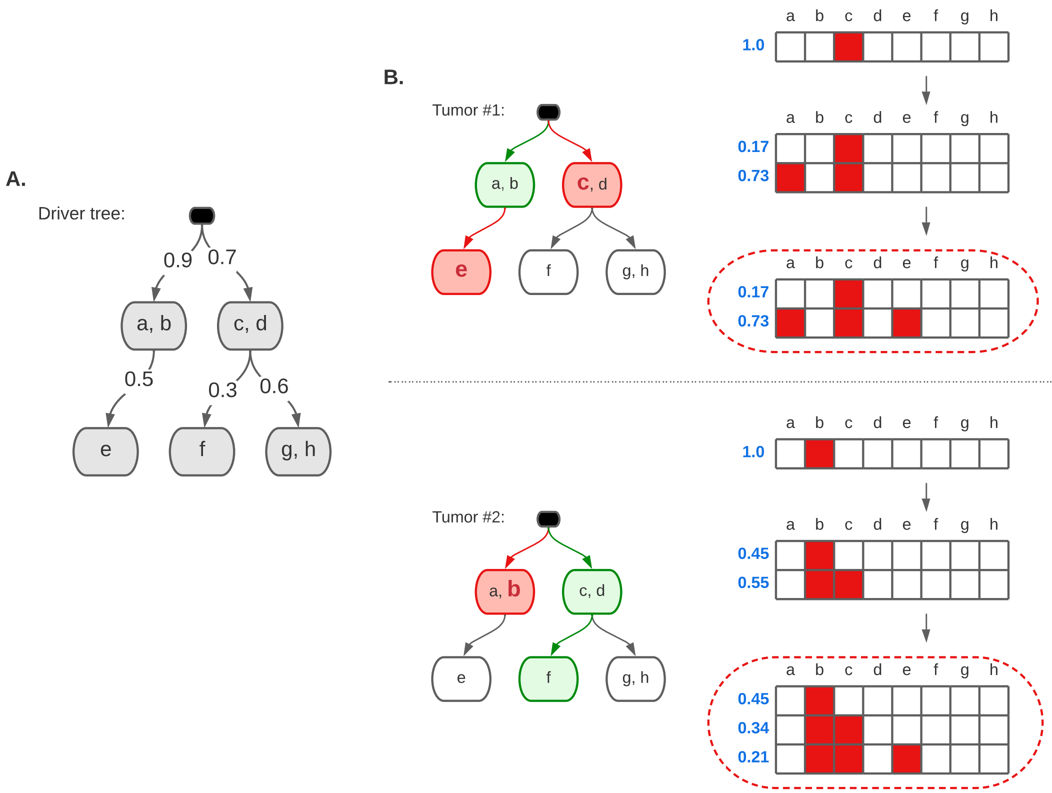

Figure 1: **A.** An example progression model. **B.** Two samples of clonal matrices corresponding to different soft decision edges (shown in green)

comprehension. In tumor 1, we have drawn 0.73 from $\text{Beta}(9, 1)$. In tumor 2, we have first drawn 0.55 from $\text{Beta}(7, 3)$. Later, we have drawn 0.38 from $\text{Beta}(3, 7)$, leading to a relative population equal to $0.38 * 0.55 = 0.21$ for the last row.

#### 3.2 Random tree structures for synthetic data experiments

To investigate our performance in recovering the generative progression models, we constructed a collection of 100 random progression models using the procedure explained in Alg. 2. Following this procedure, we get a set of progression models with an average of 1 to 4 genes per node. The average depth of the genes is also forced to be between 1 to 4 (with the root node in depth 0). Fig. 2 shows a scatter plot of our progression model features, where each point represents one of our randomly generated structures. The progression models corresponding to the red points in this scatter plot are selected as representative examples and are depicted in Fig. 3

Using each progression model in the collection, we constructed a set of 12 datasets with the number of tumors in $\{10, 30, 100, 300\}$ and the error probability in $\{0.001, 0.01, 0.02\}$, leading to a set of 1200 synthetic datasets in total. Fig. 4 shows our inferred progression models for the experiments with 300 tumors and the error probability of 0.001 for our 8 example progression models shown in Fig. 3. As shown in Fig. 4, even for the cases where we have low F-scores, e.g., cases 3 and 5, the weighted F-scores are quite high. This implies that our inference algorithm accurately recovers the relationships among pairs of highly mutated genes, which are naturally more important. We emphasize that in all these experiments, our posteriors are significantly better than the generative models, as expected.

**Algorithm 2** Construction of random generative progression models

Sample the number of genes $N$ uniformly between 10 and 40.
Sample the number of nodes $K$ uniformly between $\lceil N/4 \rceil$ and $N$. ▷ $N/K \leq 4$
Sample the probability of linear expansion $\mu$ from $\mathcal{U}(0, 1)$.
Let the average depth of the genes $\kappa = \infty$
**while** $\kappa > 4$ **do**
  Initialize the set of nodes $\mathcal{S}$ to include only the empty root node
  **for** $i \in \{1, \ldots, K\}$ **do**
    Create node $i$ including gene $i$ ▷ Each node should have at least one gene
    Sample the firing probability of node $i$ from $\mathcal{U}(0.05, 0.95)$.
    **if** Bernoulli($\mu$) **then**
      The parent of node $i$ is node $i - 1$
    **else**
      The parent of node $i$ is selected uniformly among the nodes in $\mathcal{S}$
    Add node $i$ to $\mathcal{S}$
  **for** $i \in \{K + 1, \ldots, N\}$ **do**
    Add gene $i$ to node $j$, selected uniformly from the set of nodes (except the root).
  Calculate the average depth of the genes $\kappa$
**for** $v \in \mathcal{S}$ **do**
  **if** $|D_v| > 1$ **then**
    Sample $\overrightarrow{\alpha}_v \sim$ Dirichlet(ones($|D_v|$))
Calculate the mutation rate $\alpha_g$ for all genes ▷ To be used for generating datasets

#### 3.3 Computational complexity analysis

Due to the linearity of our likelihood calculation procedure with respect to the dataset size (number of tumors $\times$ number of genes), we expect the runtime of the experiments to be linear in the dataset size. Fig. 5 shows the scatter plot of our runtime for all the 1200 synthetic data experiments and a linear regressor fitted to the points.

#### 3.4 Alternative measures for the performance

As an alternative to the F-scores we use for measuring our performance in the synthetic data experiments, we have also calculated two metrics defined in [1] for comparing hierarchical structures. Fig. 6 show the box plots for DISC and CASET distance of the inferred models to their corresponding generative models. As shown in this figure, by increasing the number of tumors or decreasing the error rate, we output a progression model which more closely resembles the model that is used to generate the data.

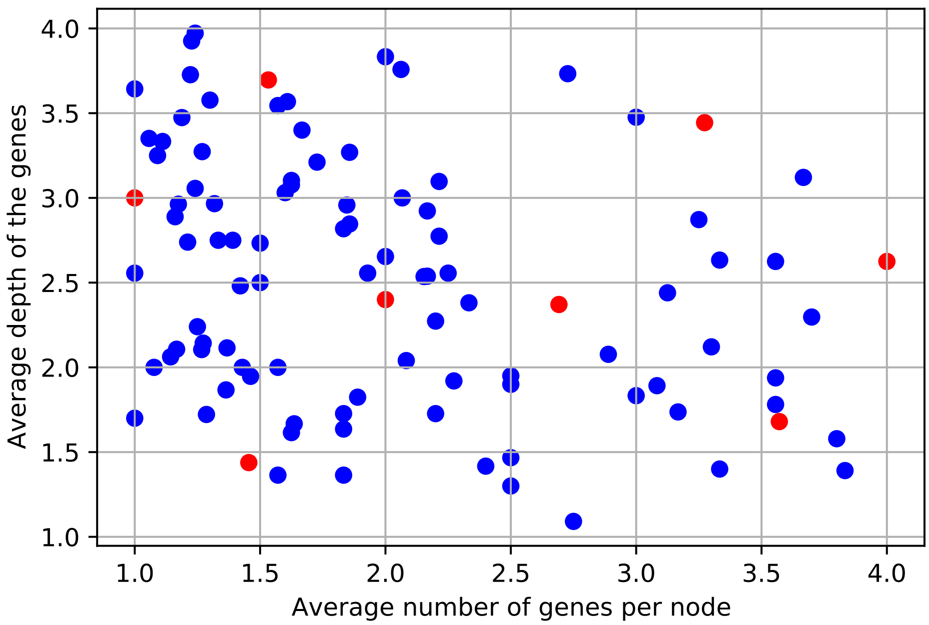

Figure 2: The average depth of the genes and the average number of genes per node in our 100 random progression models. The models corresponding to the red points are plotted in Fig. 3 as examples.

### 4 Details on the biological data analysis

#### 4.1 MI and ME matrices

To better describe the biological datasets we studied, in this section, we present the mutual inclusivity scores [2] and our MI and ME matrices that are computed to guide our MCMC algorithm. Fig. 7 shows the mutual inclusivity scores for the TRACERx LUAD dataset. Note that the MI score ranges between $-1$ and 1. The MI score of $-1$ implies perfect mutual exclusivity, meaning that no clone has mutations in both genes. On the other hand, the MI score being 1 implies perfect mutual inclusivity, meaning that the pair of genes are always mutated together. Figs. 8 and 9 show the MI and ME matrices we have computed for the LUAD dataset. These matrices keep the statistical significance of the mutual inclusivity (MI) and the mutual exclusivity (ME) signals among all pairs of genes in terms of log p-values. Note that while a broad set of mutual inclusivity and exclusivity patterns exist in the data (see Fig. 7), only a small subset of the signals are statistically significant. Figs. 10, 11 and 11 show the mutual inclusivity scores, the MI matrix and the ME matrix for the TRACERx LUSC dataset, respectively.

#### 4.2 Posterior distributions

Using an MCMC approach for making inferences, we can collect samples from the posterior distribution of the progression models, given the input data. As the paper explains, we use the maximum a posteriori (MAP) sample as our inferred model. In this section, we present our analysis of the actual posterior distribution and discuss the level of the posterior concentration around the reported best sample.

To characterize the posterior distribution, we introduce a way of encoding the structures using a binary matrix, as follows. Given a progression model including $N$ genes, we define a $N$ by $N$

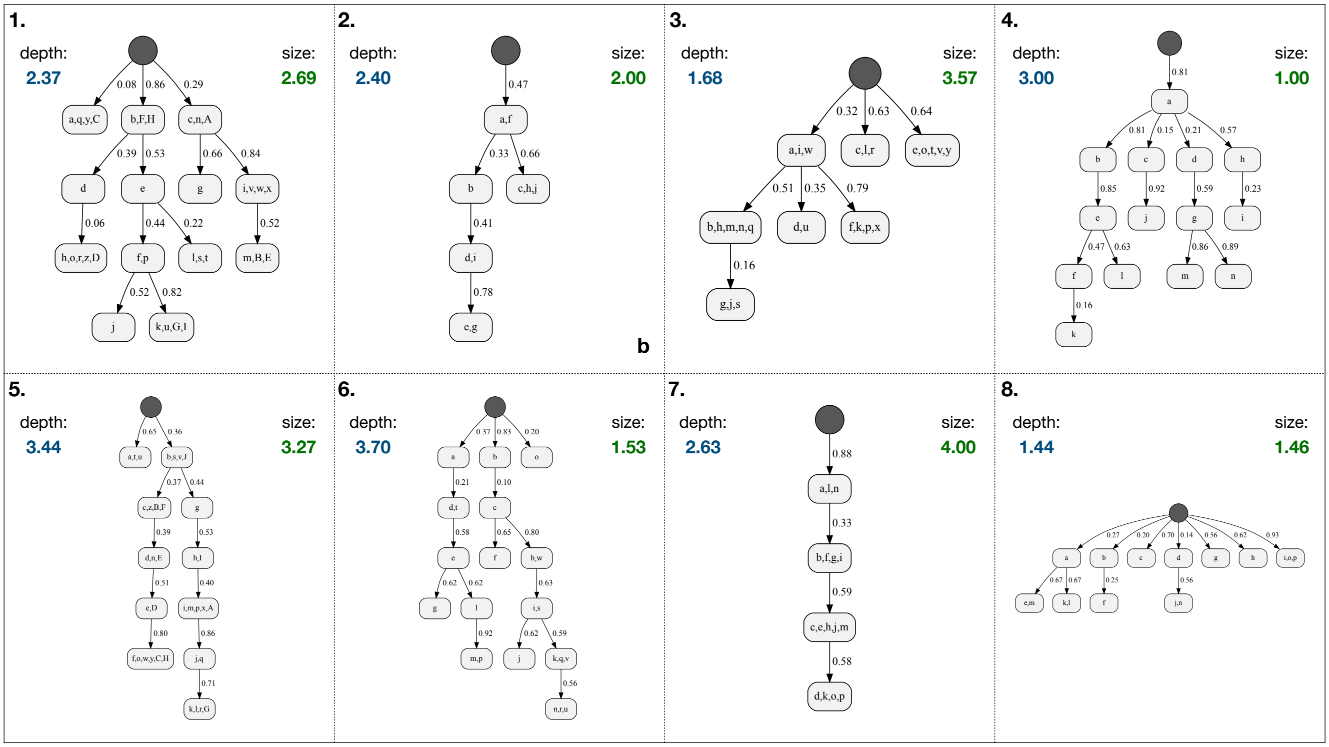

Figure 3: Example progression models with various parameters used in our synthetic data experiments. The numbers shown in blue and green on the top-left and top-right sides of each structure show the average depth of the genes and the average number of genes per node, respectively.

binary matrix to represent the structure, such that the element in the $i^{\text{th}}$ row and $j^{\text{th}}$ column is 1, if and only if the node containing gene $i$ is the direct parent of the node containing gene $j$. Taking the average of such matrices corresponding to a set of posterior samples results in a matrix with elements between 0 and 1, which can be seen as a representative of the posterior distribution. We call this matrix our posterior matrix. Given a particular sample and a posterior matrix, we can review each parent-child relation (among the gene pairs) in the sample and check its corresponding value in the posterior matrix.

We used a total of 10 chains and 100k samples to analyze each of our biological datasets. As the chains seem to have converged after 30% of the iterations, we discard the first 30k iterations as the burn-in samples and use a thinning interval of 10 to collect a total of 70k samples for computing the posterior matrices. Figs. 13 and 14 show our MAP samples beside the posterior matrices for our analysis of the TRACERx LUAD and LUSC datasets. Consider Fig. 13-B, for instance. The element in the ARID2 row and KDR column being 0.1 means that in around 10% of our samples from the posterior, the node containing ARID2 is the parent of the node containing KDR. We have annotated the relations presented in the MAP samples by the orange boxes in the posterior matrix. As shown in Fig. 13-B, most of the relations implied by the MAP sample for our LUAD analysis are present in most of our posterior samples. For the LUSC results, on the other hand, as shown in Fig. 14-B, the posterior distribution seems to be smoother around the MAP sample, and there are some highly repeated relations like TP53 to FAT1 that are not present in the MAP sample. Interestingly, the TP53 to FAT1 relation is the only relation present in the MHN network we got from our TreeMHN analysis of the LUSC dataset, as shown in the paper. A 60% majority of our posterior samples imply this relationship as well.

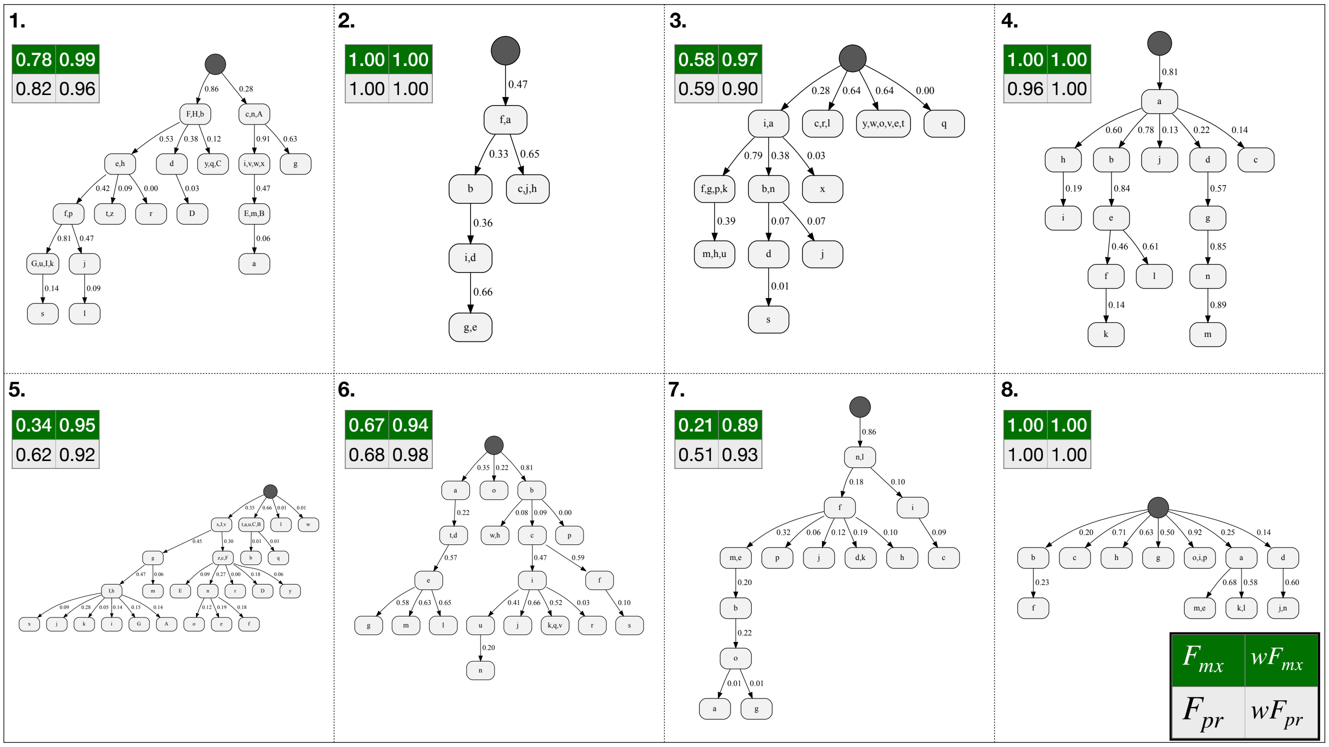

Figure 4: Our inferred progression models (MAP samples) for the experiments with the progression models shown in Fig. 3 with 300 tumors and the error probability of 0.001. The F-scores (normal and weighted versions) for identification of the mutual exclusivity and progression relations are shown beside each model. See the legend box on the bottom right side of the figure.

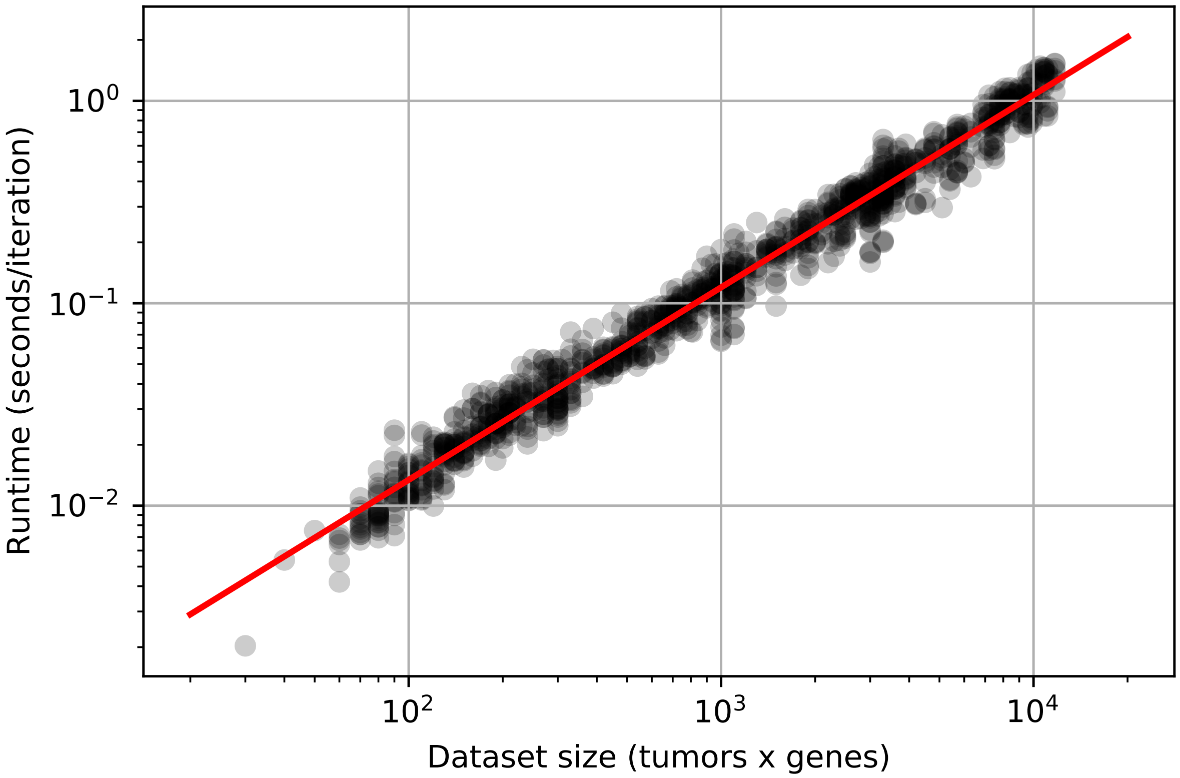

Figure 5: Runtime of the inference algorithm with respect to the dataset size. The slope of the linear regressor is 0.95 (note the log-log scale).

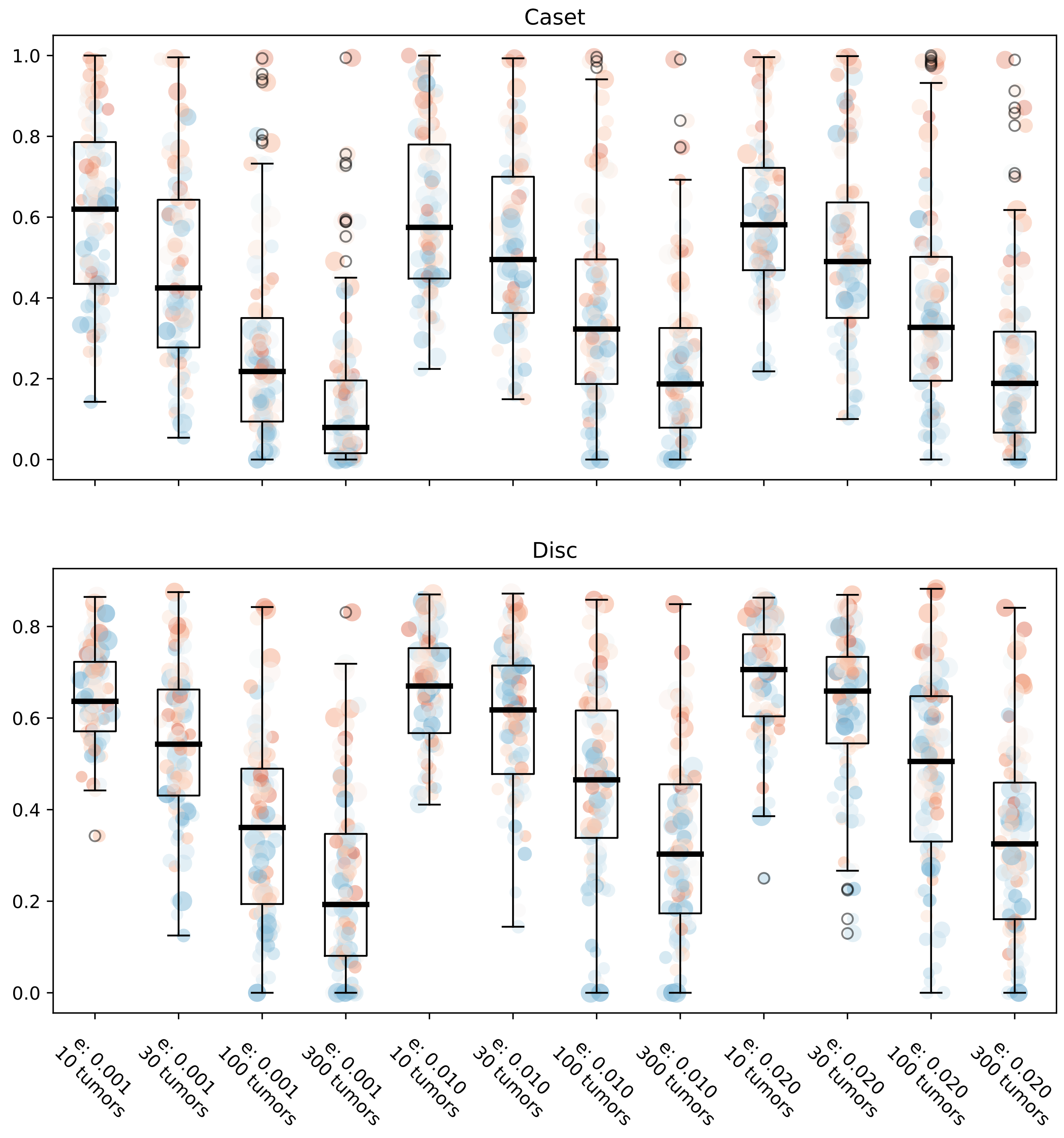

Figure 6: DISC and CASET distance between the inferred models and their corresponding generative models. Both DISC and CASET take values between 0 and 1, with lower distance implying more similarity.

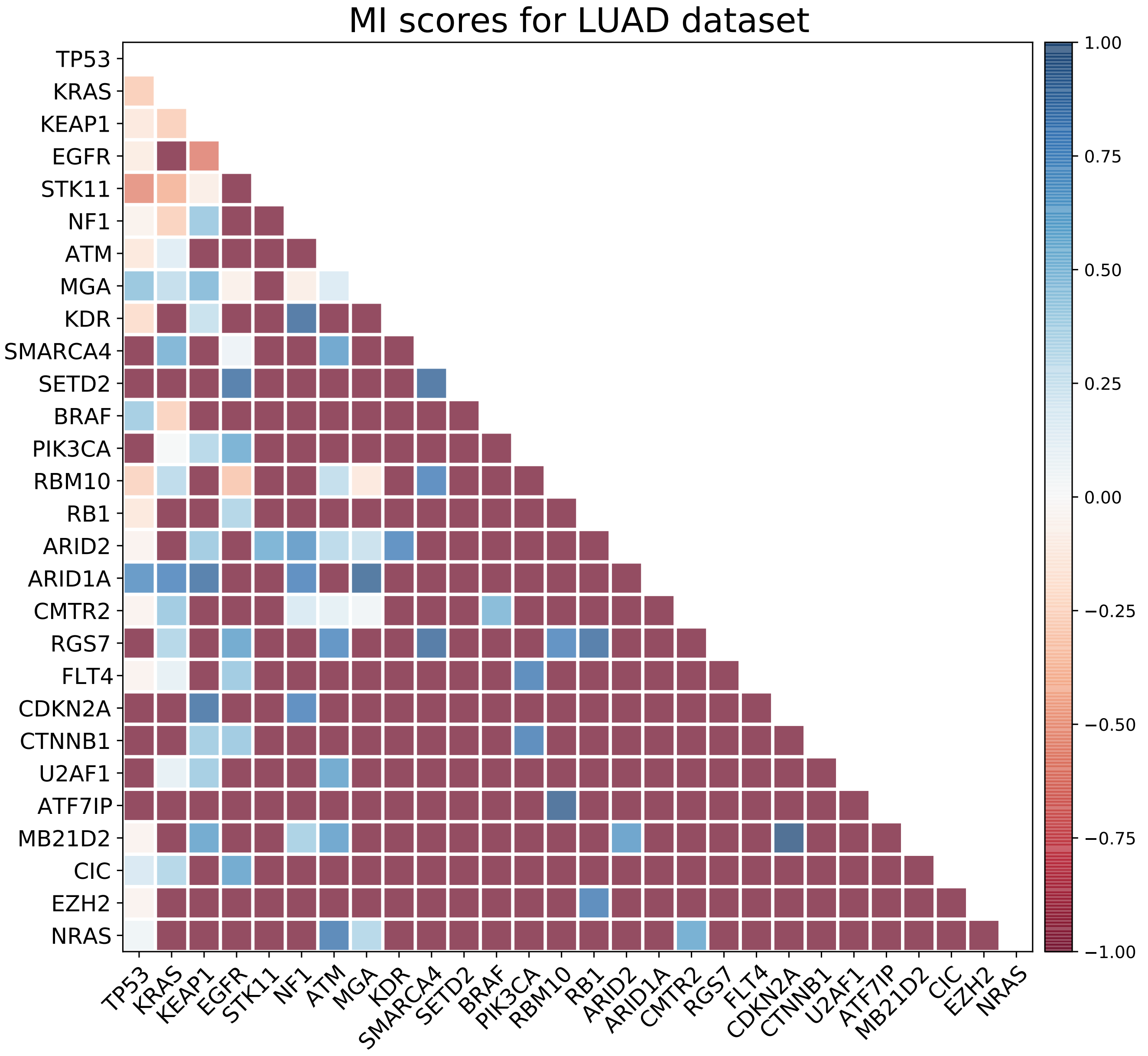

Figure 7: The mutual inclusivity (MI) scores computed for the LUAD dataset.

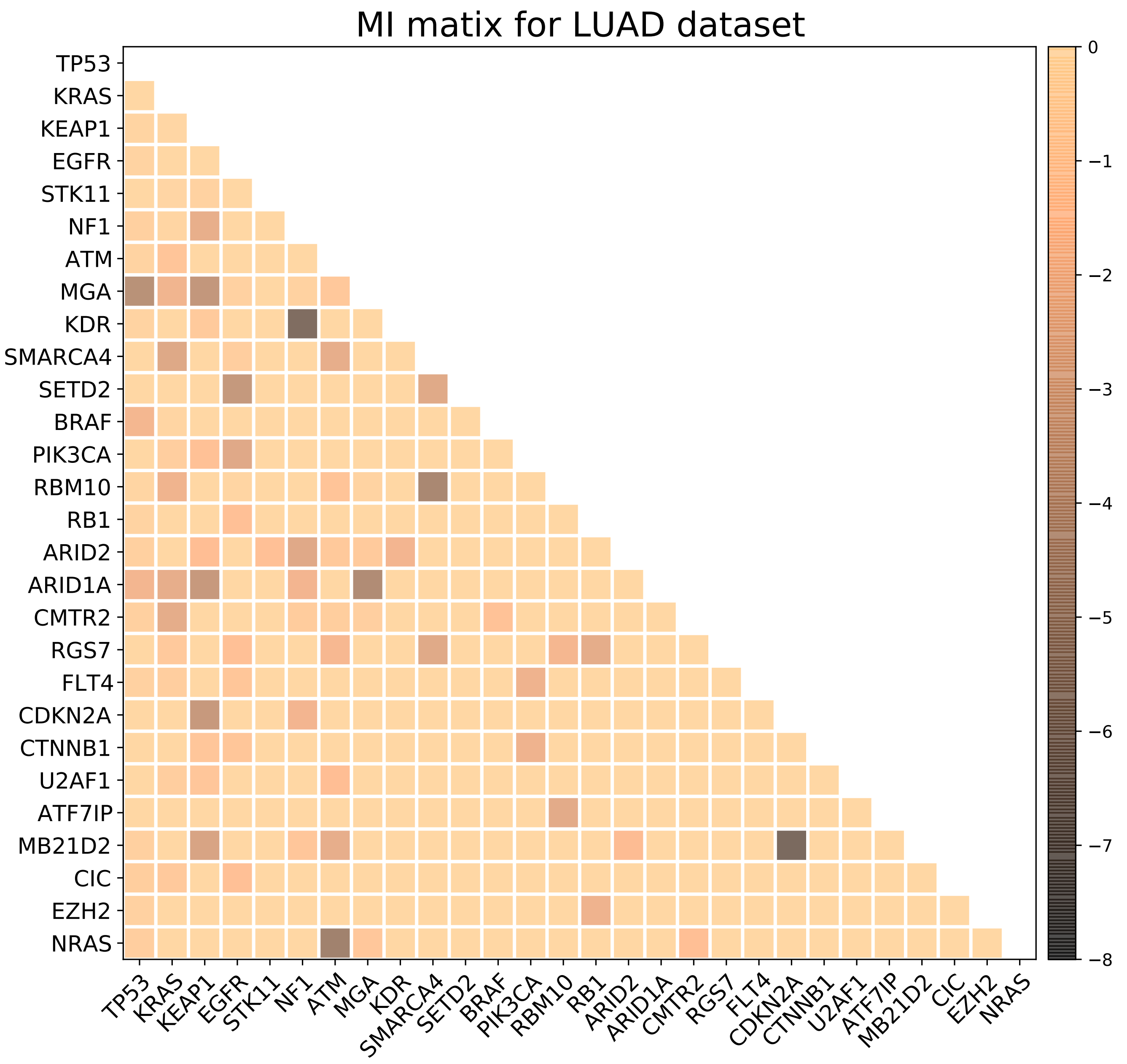

Figure 8: The MI matrix for the LUAD dataset.

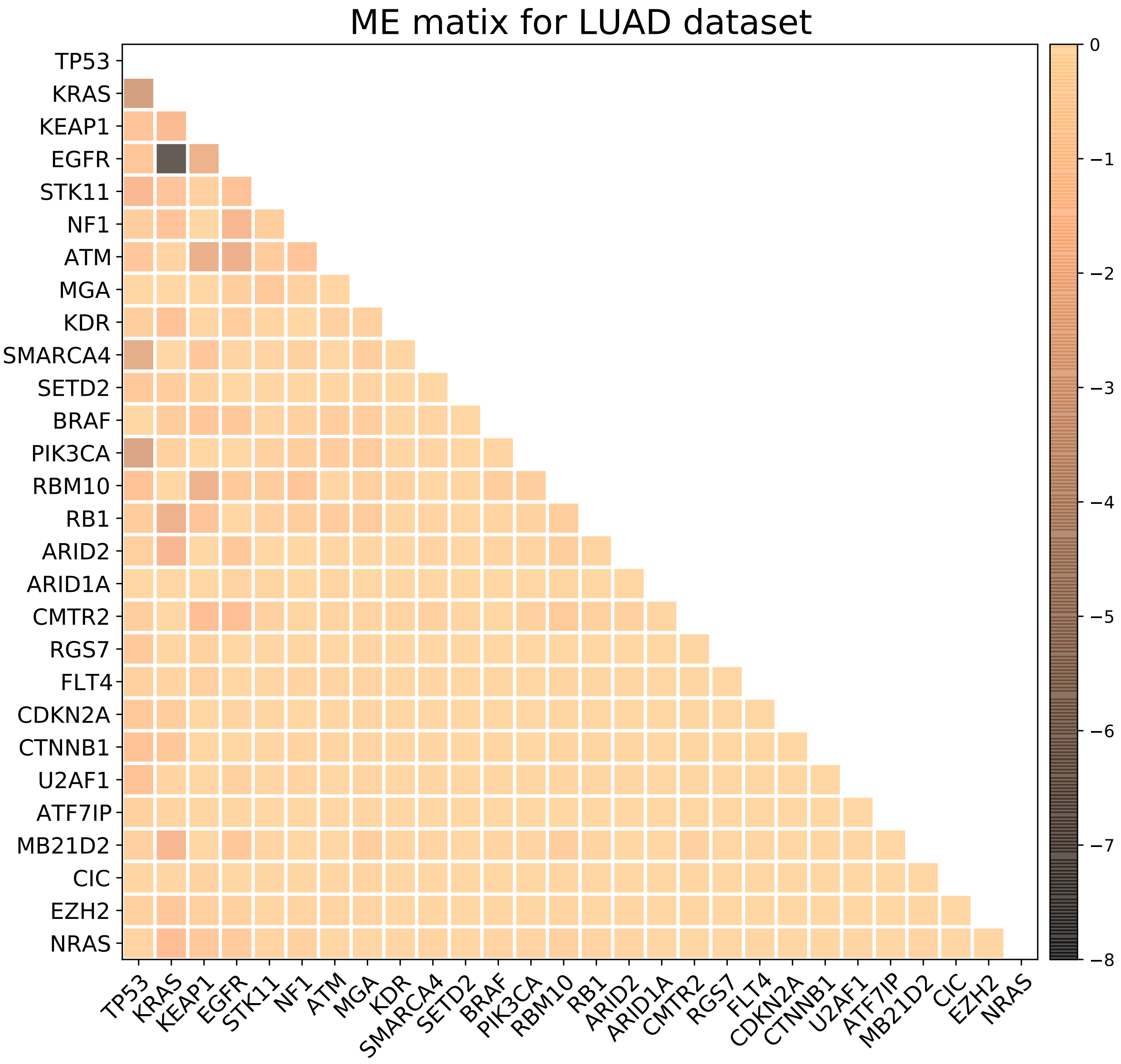

Figure 9: The ME matrix for the LUAD dataset.

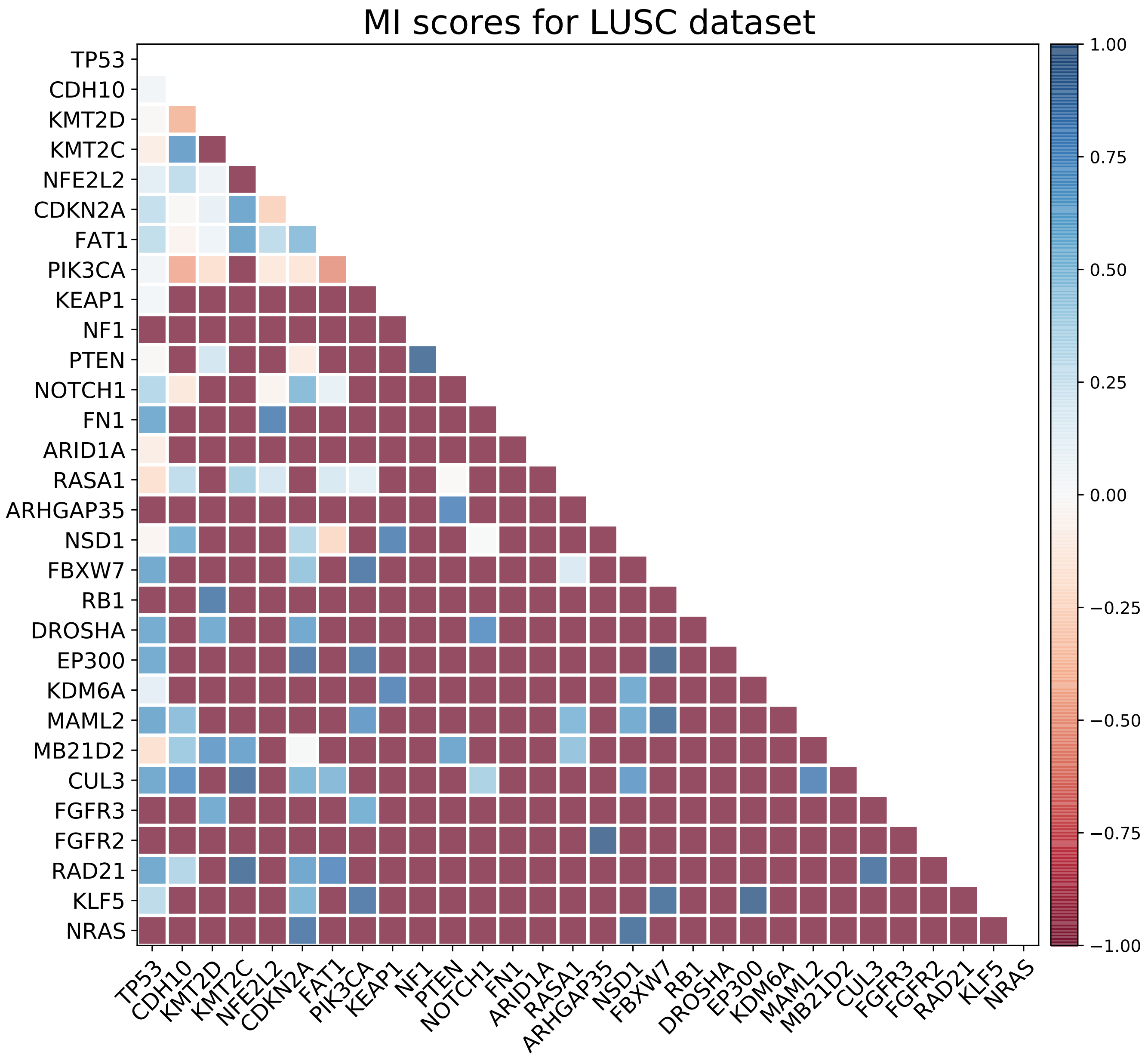

Figure 10: The mutual inclusivity (MI) scores computed for the LUSC dataset.

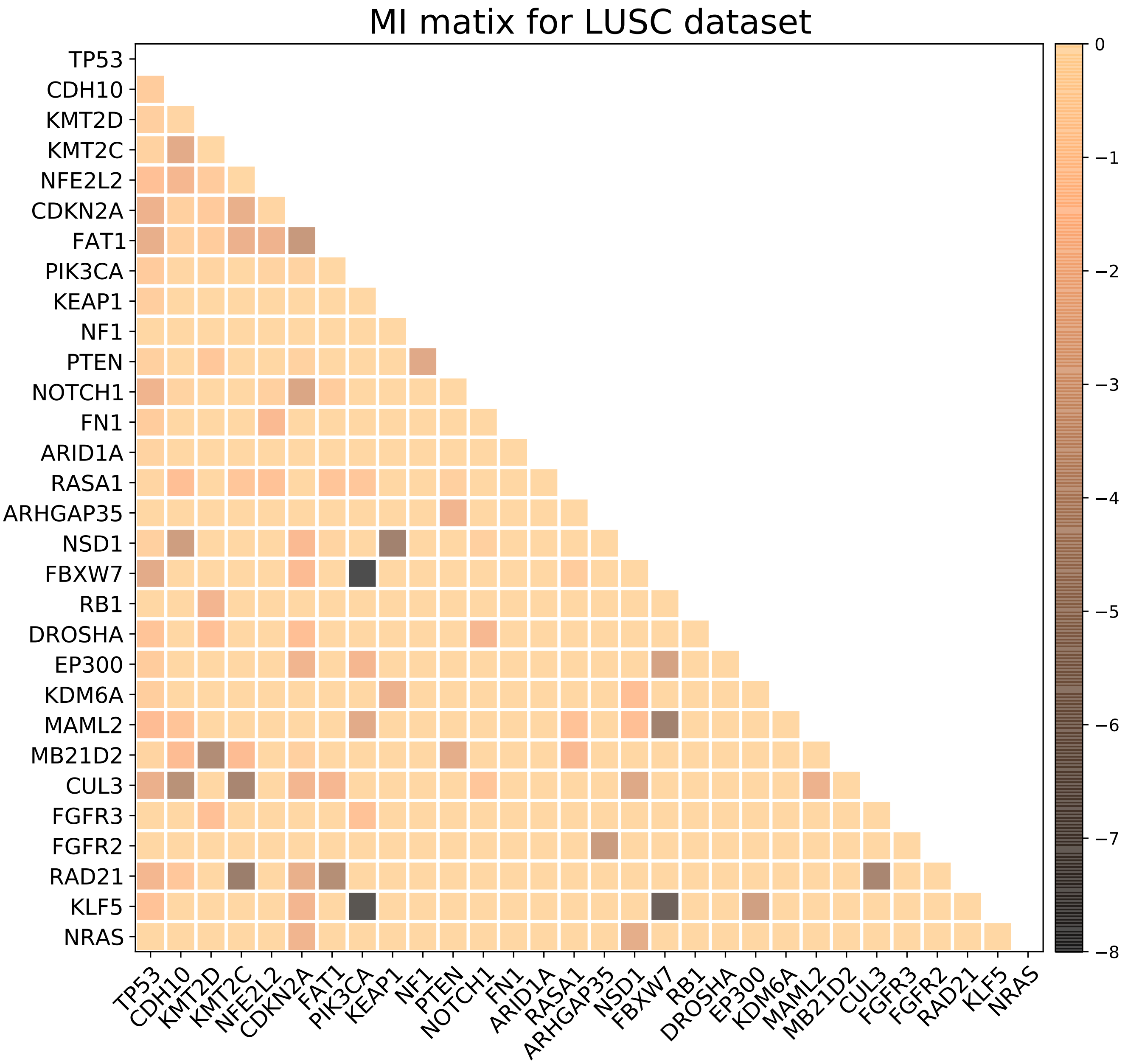

Figure 11: The MI matrix for the LUSC dataset.

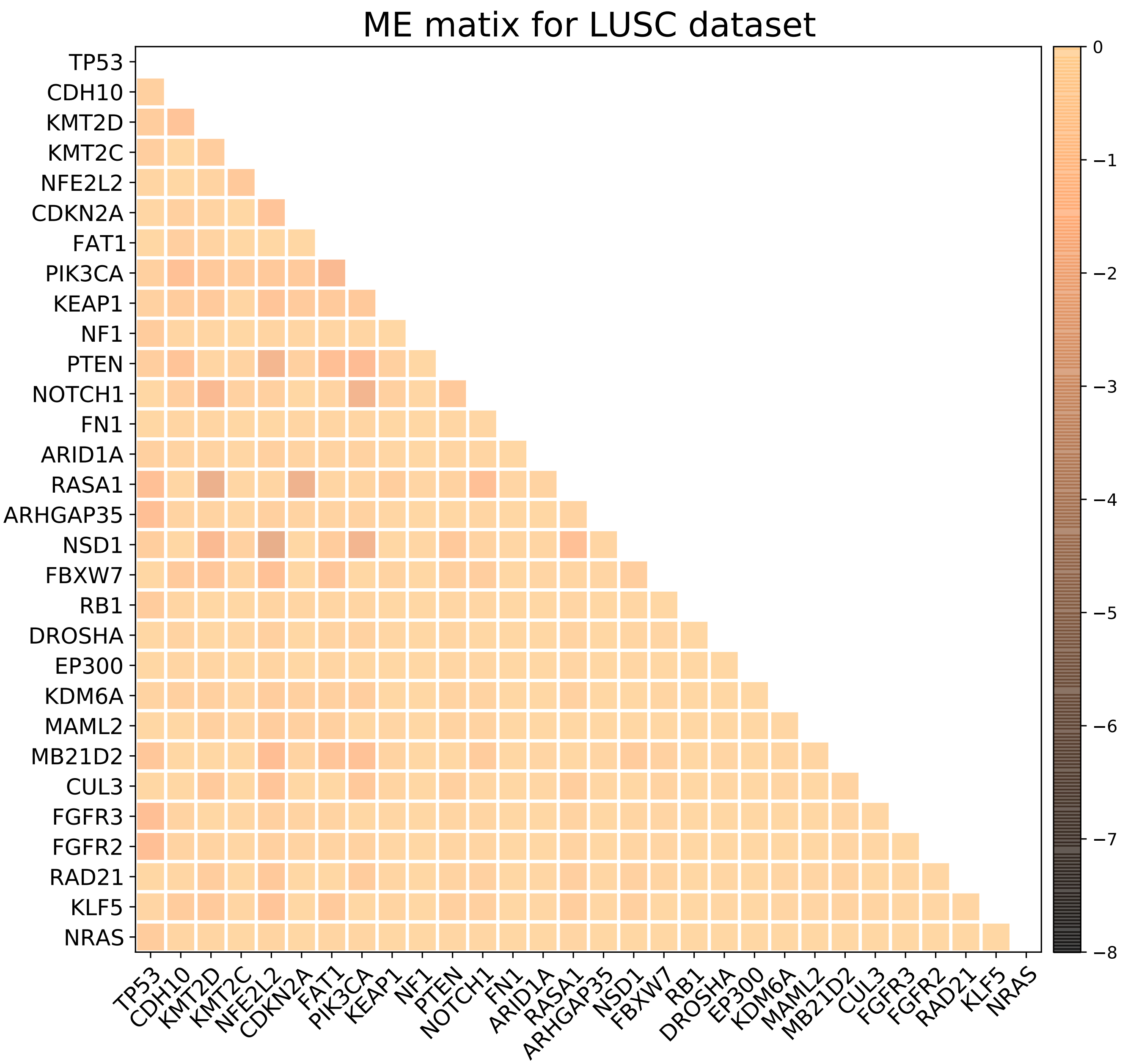

Figure 12: The ME matrix for the LUSC dataset.

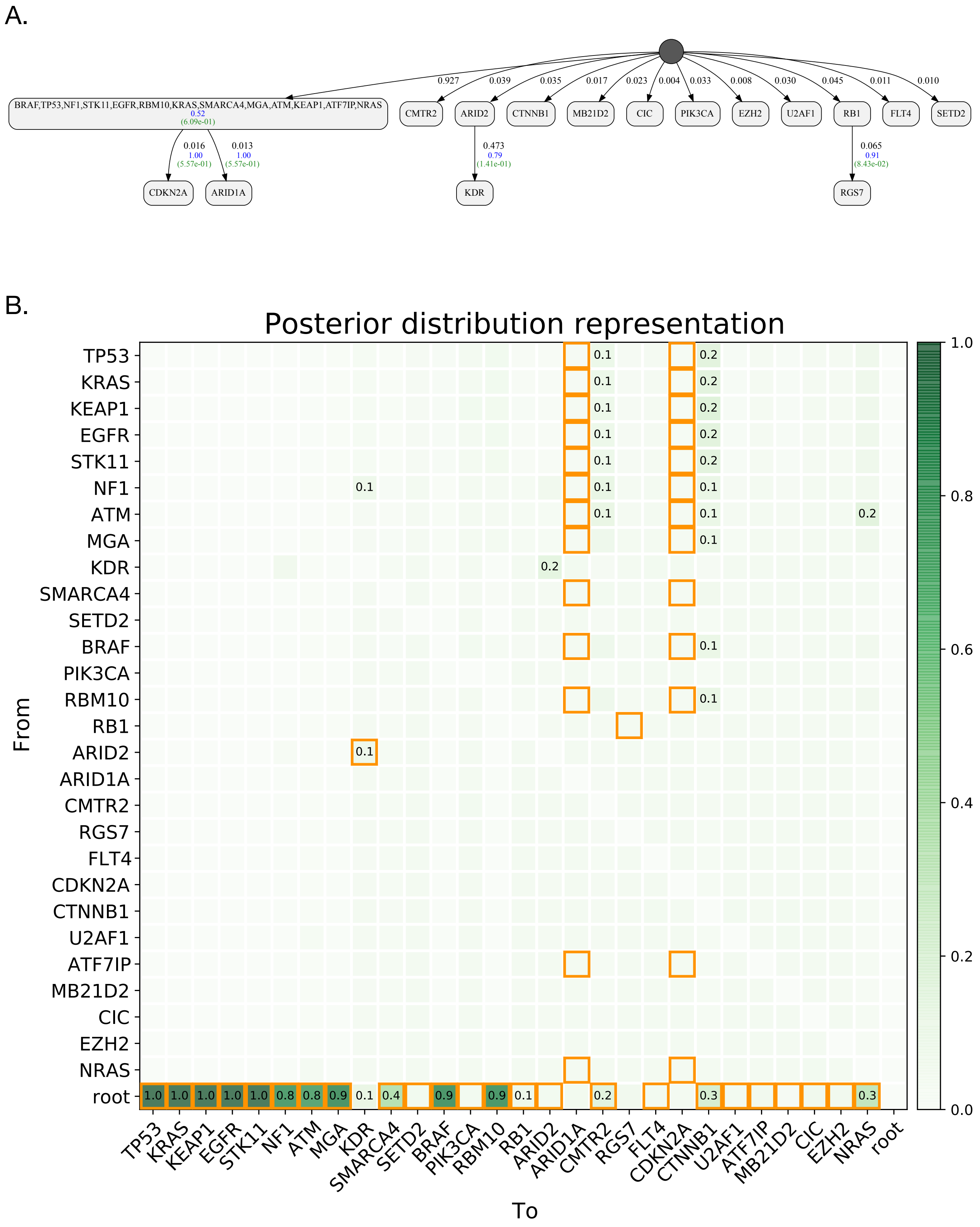

Figure 13: Results of our TRACERx LUAD analysis. **A:** The MAP sample, reported as the inferred model. **B:** The posterior matrix, where the elements corresponding to the MAP sample are annotated with the orange boxes.

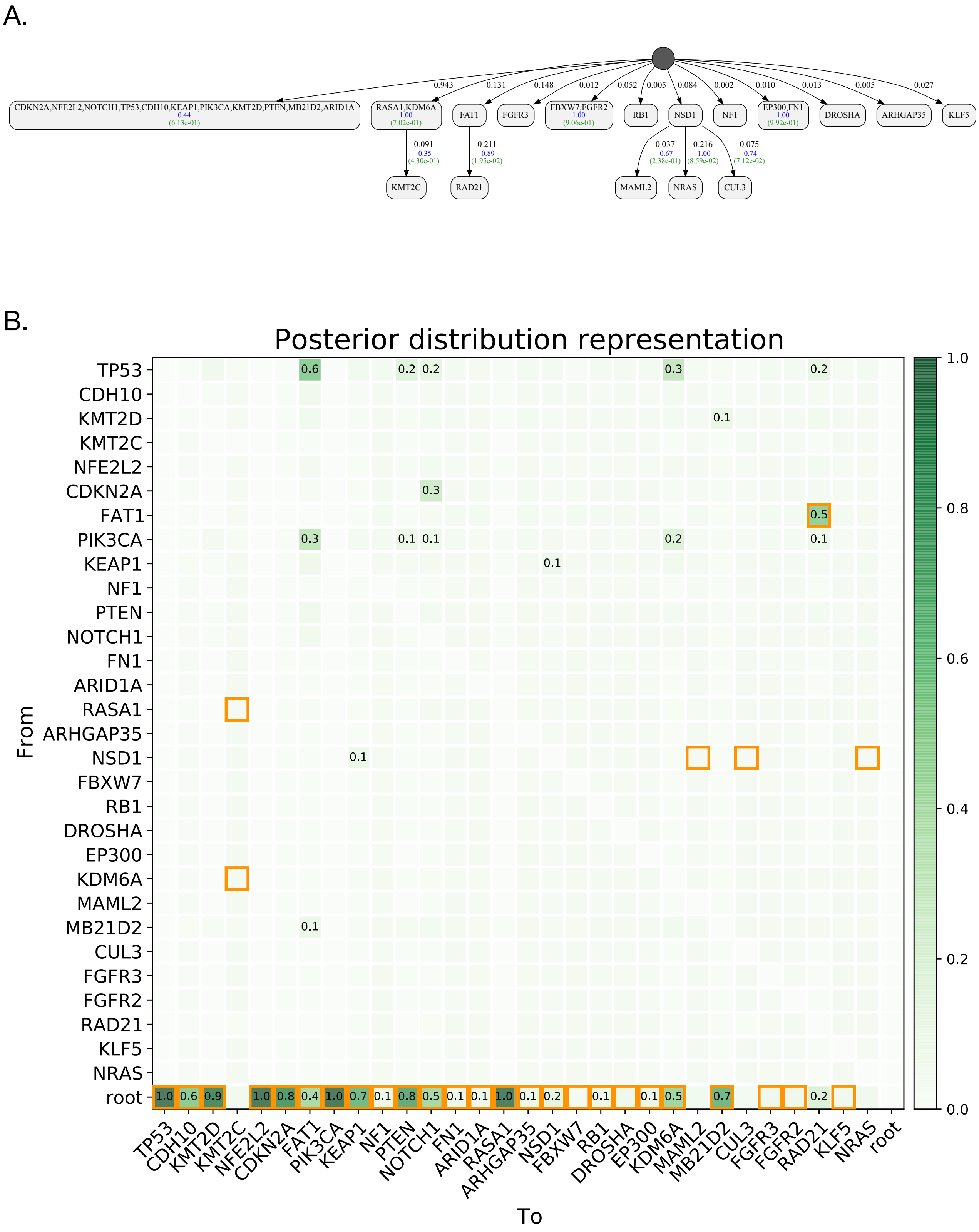

Figure 14: Results of our TRACERx LUSC analysis. **A:** The MAP sample, reported as the inferred model. **B:** The posterior matrix, where the elements corresponding to the MAP sample are annotated with the orange boxes.
